# Neural correlates of object identity and reward outcome in the corticohippocampal hierarchy: double dissociation between perirhinal and secondary visual cortex

**DOI:** 10.1101/2023.05.24.542117

**Authors:** J. Fiorilli, P. Marchesi, T. Ruikes, G. Huis in ‘t Veld, R. Buckton, M. Duque Quintero, I. Reiten, J. Bjaalie, C.M.A. Pennartz

**Affiliations:** Cognitive and Systems Neuroscience Group, Swammerdam Institute for Life Sciences, Faculty of Science, University of Amsterdam, Amsterdam, Netherlands; Department of Molecular Medicine, Institute of Basic Medical Sciences, University of Oslo, Norway

## Abstract

Neural circuits support behavioral adaptations by integrating sensory and motor information with reward and error-driven learning signals, but it remains poorly understood how these signals are distributed across different levels of the corticohippocampal hierarchy. We trained rats on a multisensory object-recognition task and compared visual and tactile responses of simultaneously recorded neuronal ensembles in somatosensory cortex, secondary visual cortex, perirhinal cortex and hippocampus. The sensory regions primarily represented unisensory information, while hippocampus was modulated by both vision and touch. Surprisingly, secondary visual cortex but not perirhinal neurons coded object-specific information, whereas perirhinal but not visual cortical neurons signaled trial outcome. A majority of outcome-related perirhinal cells responded to a negative outcome (reward omission), whereas a minority of other cells coded positive outcome (reward delivery). Our results support a distributed neural coding of multisensory variables in the cortico-hippocampal hierarchy, with a double dissociation between higher visual cortex and perirhinal cortex in coding of object identity versus feedback on trial outcome.

## Introduction

Object recognition serves animals to guide themselves to goals of interest, such as locations with food. Multisensory object representations are thought to be encoded by a hierarchically organized, cortical network in which unisensory information is initially processed in distributed unimodal regions and converges in higher order areas. The perirhinal cortex (PER) is often considered to be a central hub in this hierarchy, because of its reciprocal anatomical connections with, among others, the temporal association cortex, somatosensory whisker system and visual cortex (Murray and Bussey 1999; Eichenbaum 2000; Agster & Burwell, 2009; Jacklin *et al.,* 2016; Alvarez & Squire 1994; Fiorilli et al. 2021). Lesion studies in rats that spontaneously explored their environment demonstrated that PER damage causes impairments in zero-delay recognition for objects having complex visual feature conjunctions and in crossmodal object recognition (Bartko *et al*., 2007; Albasser *et al*., 2010; Reid, Jacklin, & Winters, 2012). As a first hypothesis to be considered here, it has been proposed that PER contributes to object (or item) perception in addition to memory (Dickerson and Eichenbaum 2010), especially when multiple object features or modalities have to be processed. Despite the lesion evidence, no electrophysiological studies have thus far compared the influence of sensory inputs during object recognition between the PER, the sensory neocortical regions and hippocampus of rodents (HPC).

A second hypothesis on the functionality of PER implies this structure in the coding of task and reward contingencies by interacting with interconnected structures involved in planning, motivation and affect, such as the medial prefrontal cortex (mPFC), orbitofrontal cortex and amygdala (Agster *et al*., 2016, Kajiwara *et al*., 2003; Burwell, Witter & Amaral, 1995; Fiorilli *et al*. 2021). Studies in monkeys showed that the PER carries information about cued reward schedules rather than the physical properties of sensory cues themselves, such as brightness (Eradath *et al*., 2015; Liu and Richmond, 2000). In rats, PER neurons have been shown to code for spatial segments of a task environment (Bos *et al*., 2017), but also to represent behavioral choices on a fine time-scale during spatial responses on a touch screen (Ahn & Lee 2015). Spatial correlates in PER are generally not reported in the absence of spatial task constraints (e.g. when foraging in an open field arena; Burke *et al*., 2012; Deshmukh *et al*., 2012), suggesting that spatial-navigational correlates in PER may arise only when paired with reward prediction. Despite the available evidence for a contribution of PER to multisensory processing and value-based coding, it remains debatable whether sensory and value-based representations are both present in PER, and how PER compares to other regions in the cortico-hippocampal hierarchy.

The current study aimed to quantify sensory-evoked (tactile and visual) firing responses in the PER during multisensory object sampling, as well as firing-rate modulation by choice behavior and reward delivery. We additionally set out to compare these neural representations in PER to those simultaneously recorded from other regions of the cortico-hippocampal system. We devised a reward-driven multisensory object recognition task in which rats repeatedly discriminated between two familiar, solid 3D objects via whisker touch and/or vision. Object sampling enabled the rats to make a spatial choice to approach one of two goal locations, each of which was associated with one of the objects and yielded reward when correctly chosen. During this task, we simultaneously recorded ensemble activity with single-cell resolution from primary somatosensory barrel cortex (S1BF), secondary visual cortex (V2L), PER and HPC (including CA1, CA3 and DG).

Our results revealed distinct functional contributions of V2L and PER within the cortico-hippocampal hierarchy. First, V2L neurons carried information about object identity during visual sampling, but specific object representations were neither found in PER nor HPC. We additionally report a much higher sensitivity to tactile and visual inputs for HPC and the sensory cortical regions compared to PER. Second, representations in PER maximally differentiated between the choice sides (left vs. right) when rats kept their snout poked into one of two reward ports, indicating a specific role in coding of expected reward for a given goal location. Third, whereas modulations by trial outcome in V2L (and in S1BF and HPC as well) correlated to different post-reward behaviors (reward consumption vs. exiting the reward port), outcome-related responses in PER occurred earlier, viz. when rats started to sample for reward delivery by licking, and were mainly driven by trial outcome (i.e., reward delivery or omission). Collectively, these results reveal a double dissociation between at least two of the four recorded areas. V2L encodes object identity in visual trials and is therefore likely involved in representing visual features during object discrimination. In contrast, PER cells anticipated arrivals at a given goal location and represented unexpected trial outcome, suggesting a prominent contribution of PER in encoding motivational value of events, rather than a general role in object perception and recognition.

## Results

### Behavioral results and multi-area electrophysiology

We trained freely behaving rats (N=4) on a multisensory two-object recognition task, in which solid objects had to be associated with different choice sides in order to obtain reward. During a given trial, object sampling was either possible by whisker palpation, vision, or by both senses combined (Fig. 1a). After an intertrial interval (ITI; 12 seconds) rats could sample an object, located a gap distance away from the elevated platform. Whisker palpation was prevented in visual trials by having a larger gap distance to the object, and availability of visual object information was controlled with an object-focused illumination in an otherwise darkened room. This illumination was triggered when rats perched across the gap and towards the object, forcing a similar pose and head-position across the different trial types.

**Figure 1:**
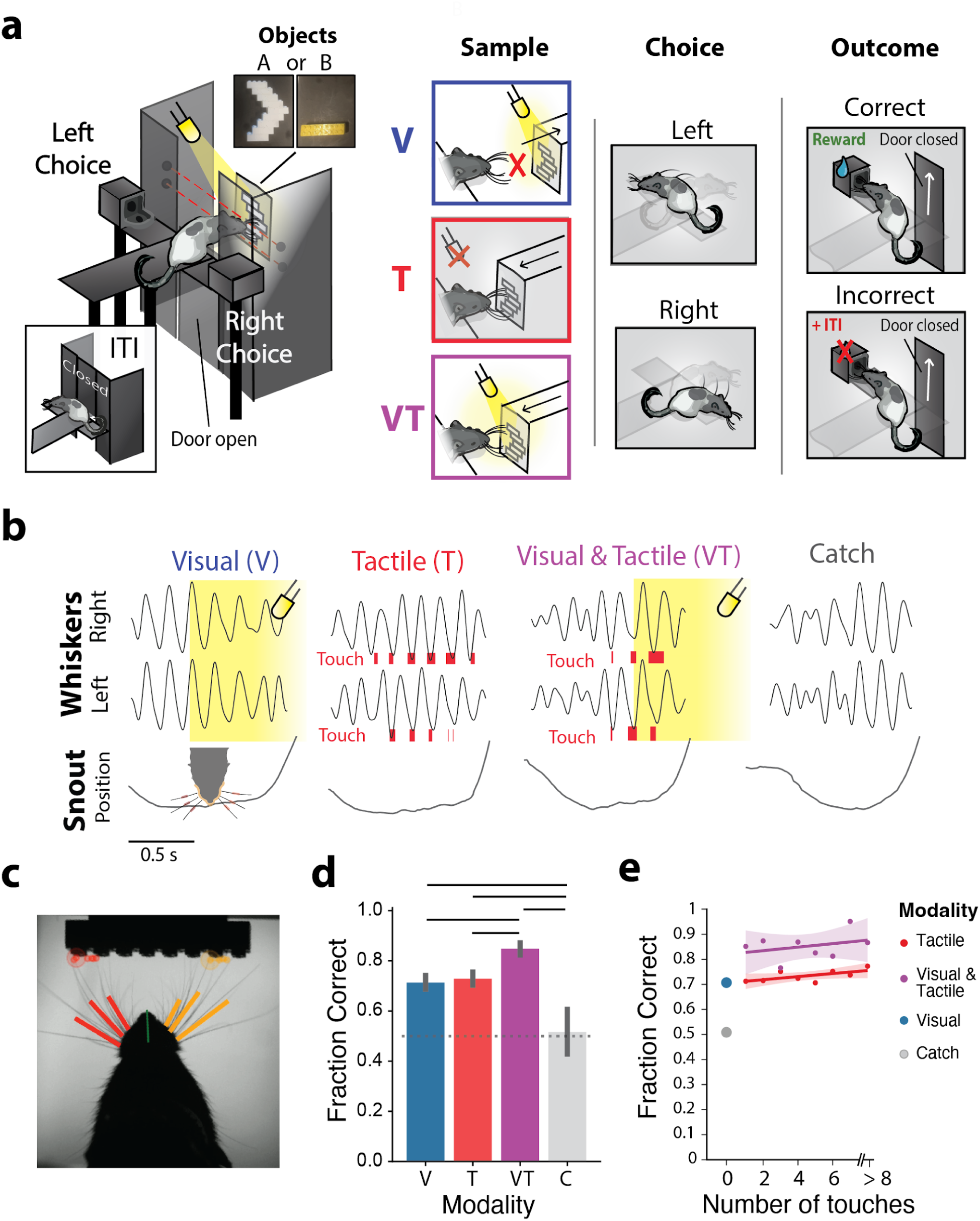
Performance and single trial examples of the multisensory object discrimination task. **a)** Schematic overview of the reward-driven discrimination task. Rats were trained to associate two objects with opposed reward sides. Different trial types allowed object sampling either by vision (V), touch (T), or by both senses combined (VT). **b)** Example trials with object approach and time course of whisking behavior. Rats started whisking during the approach towards the object. In tactile and multisensory trials, rats could reach and palpate the object with its whiskers. In visual and multisensory trials, an LED light illuminated the object when the snout was in front of the object. The lower row displays the snout distance from the object over time, during the approach to and withdrawal from the object. Touch epochs are indicated in red and illumination in yellow. **c)** Example frame taken from a high-speed video during tactile object sampling, with overlaid whisker and snout tracking. **d)** Rats performed the task above chance-level in all trial types, taking the catch trials with no object information (neither visual or tactile) as the chance-level comparison. Performance in multisensory trials was significantly higher than in visual and tactile trials (P<0.05, one-way Anova (F= 31.376, P < 0.000) with post-hoc Tukey test). Bars indicate 95% CIs. **e)** Task performance over amount of touches during object sampling. Multisensory sampling improved task performance regardless of the number of touch epochs. Lines are linear fits and shaded areas indicate the 95% CI.

Rats whisked in the air during all trials (Figure 1b; c; SI: Video), also when no tactile information was available.

The rats subsequently retracted their body from the object-sampling port and turned to make a left or right nose-poke, where the correct choice depended on object identity. Pokes on the correct object-associated side were rewarded with sucrose delivery. An incorrect response led to an extended ITI (20 s) without reward delivery. To verify that rats could not use odors or sounds related to the automated object-presentation mechanism, we also introduced catch trials. During these catch trials the object-presentation mechanism remained active so that objects were rotated into place for future presentation during the ITI, but illumination was kept off and the object itself was out of whisker reach. Rats were expected to perform around chance-level for these catch trials if they based their decision solely on visual or tactile information. Rats learned to discriminate between objects with an above-chance accuracy in all trial types, except for catch trials (Fig, 1d; Visual trials: T(27) = 14.80, P <0.001, Tactile trials: T(27) = 16.33, P =<0.001, Multisensory trials: T(27) = 26.93, P <0.001; Catch trials: T(27) = 0.34, P =1.00, one sample t-test, Bonferroni-corrected). Discrimination performance was significantly higher in multisensory than unisensory trials, indicating that rats used both their vision and somatosensory information to guide their decisions (Fig. 1d; Multisensory versus visual trials: P=0.001; Multisensory versus tactile trials: P=0.034; one-way Anova (F=31.37, P<0.001) with post-hoc Tukey test). Multisensory sampling improved task performance regardless of the number of touches (Fig. 1e).

We simultaneously recorded the activity of neuronal ensembles in multiple regions along the cortico-hippocampal hierarchy during 27 sessions from four well-trained rats. Each animal was chronically implanted with a quaddrive that contained 36 tetrodes enabling simultaneous recording of single units (SUs) from S1BF (N=91), V2L (N=279) and PER (N=191, areas 35/36, central-caudal region; Fig. 2; cf. Bos *et al*. 2017, Vinck *et al*. 2016). In two of these animals, we additionally recorded from the dorsal HPC (N=179; Subfields: CA1=59, CA3=88, DG=32). Units in the HPC were pooled across subfields for analysis, unless specified differently.

**Figure 2:**
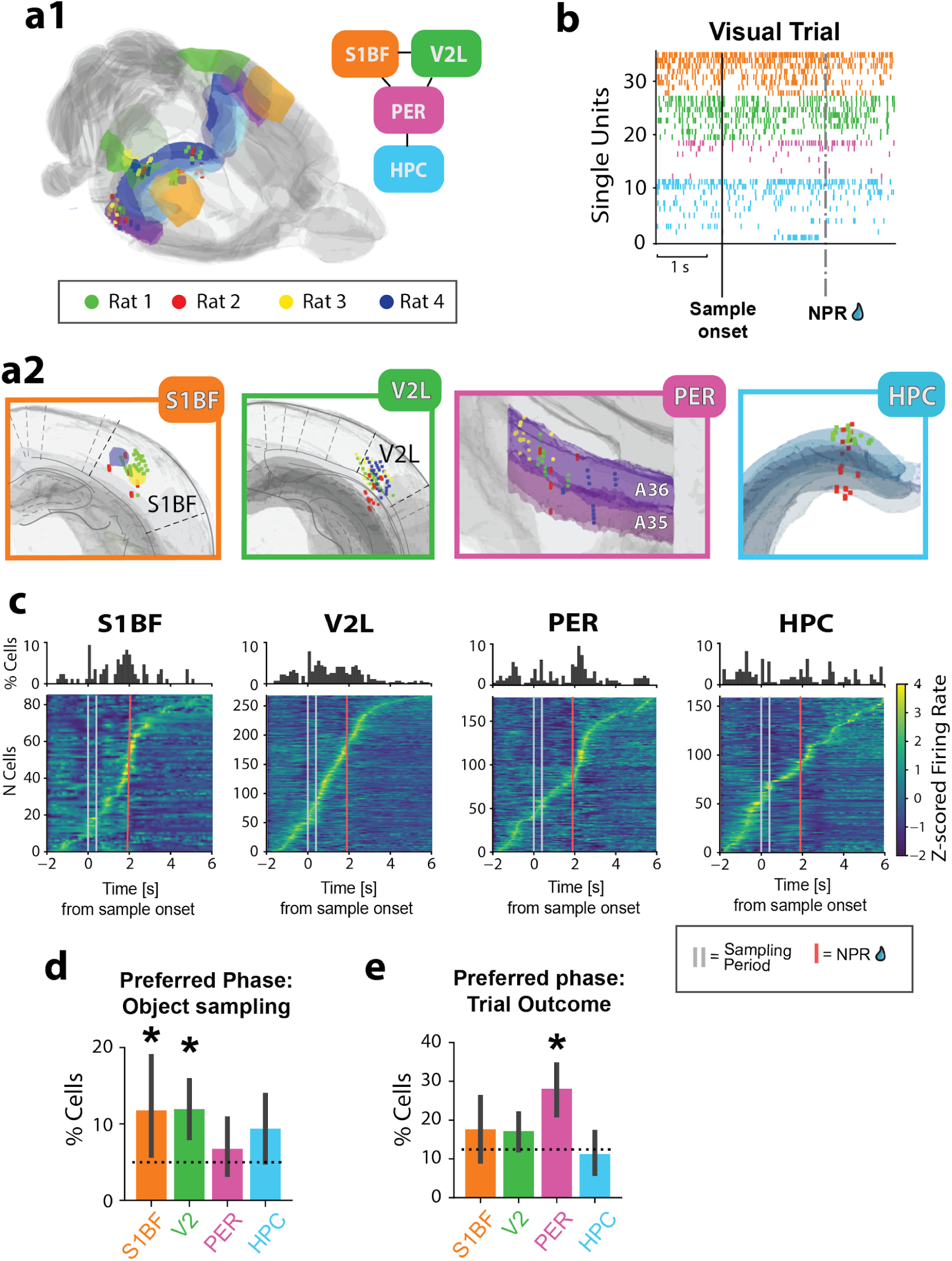
Simultaneous multi-area ensemble recordings from the cortico-hippocampal hierarchy during the object recognition and outcome phases of the task. **a1-2)** Three-dimensional spatial registration of tetrode recording locations. **a1)** Overview of all recording locations mapped on the Waxholm space atlas, color-coded per rat. Top-right schematic inset depicts the crude anatomical organization of the four recorded areas situated along the cortico-hippocampal hierarchy. **a2)** Close-up views of target regions S1BF, V2L, PER and HPC with marked recording locations. **b)** Simultaneously recorded spike trains from S1BF, V2L, PER and HPC single units during a visual trial. Units are ordered according to average firing rate and grouped per recording region. See panel (a) for color coding of brain regions. NPR is nose poke for reward. **c)** Overview of task-phase modulation in S1BF, V2L, PER, and HPC cells. The histograms indicate the percentage of cells that discharged maximally on a given time-point. The heat-maps are Z-scored firing rates of cells over time, which are linearly time-warped to account for unequal durations in object sampling and response latencies. Cells are ordered by the moment of peak firing rate relative to sample onset. **d)** Percentages of cells from a given area that preferentially discharged during object sampling. Bars are 95% CIs, dashed lines indicate chance levels given homogeneously distributed events over time. Asterisks indicate significant above-chance percentages. S1BF: 11.8; V2L: 11.9; PER: 6.7; HPC: 9.3. **e)** Percentage of cells from a given area that preferentially discharged following the NPR, regardless of reward delivery, up to 1 second after poke (S1BF: 17.6; V2L: 17.2; PER: 28.1; HPC: 11.3%). Bars indicate 95% CIs, * for significantly higher percentages than uniformly distributed events (P<0.05, two-proportions z-test)

### Tactile and visual stimulation during object sampling evokes neural activity in sensory cortices and hippocampus, but not in perirhinal cortex

The PER is thought to be richly supplied with object information from the different sensory cortices. We initially asked whether PER neurons preferentially discharge during object sampling regardless of the sensory modality, as predicted by the hypothesis that PER primarily subserves object perception. We subdivided behavioral trials into an object sampling phase, a choice phase (marked by locomotion and left vs. right turn), and a trial outcome phase initiated by a nose poke for reward (NPR). We found no preferential firing during object sampling in PER (peak firing rate probability tested against chance level of uniformly distributed events, PER probability: 0.067, Z= -3.06, P= 3.995, proportions *z*-test, Bonferroni-corrected; Fig. 2c,d; Fig. S3). Instead, PER cells preferentially discharged following the nose poke for reward (NPR; PER probability: 0.294, Z=9.79, P<0.001, proportions *z*-test, Bonferroni-corrected; Fig. 2c,e; Fig. S3). In contrast, most cells in sensory regions S1BF and V2L discharged most strongly during the choice/locomotion phase (S1BF probability: 0.40, Z=5.65, P<0.001; V2L probability: 0.39, Z=9.79, P<0.001, proportions *z*-test, Bonferroni-corrected).

Under our first hypothesis, holding that the PER contributes to complex perception by integrating multimodal object information (such that PER neurons are responsive to more than one modality), we characterized neural responses during tactile, visual, and multisensory object sampling. Using area under the receiver operating characteristic curve (AUROC) analysis, we quantified the proportion of cells significantly modulated by either light onset or whisker-object contacts.

### Visual responses

We found significant proportions of cells in area V2L and HPC that were modulated during visual object sampling (before the animals retracted from the object; Fig. 3a-d; Percentages of cells modulated by light onset in absence of simultaneous tactile sampling: V2L: 39.0%, Z=10.75, P<0.001; HPC: 13.7%, Z= 2.98, P=0.010, and during simultaneous visual and tactile sampling: V2L: 31.0%, Z=8.84, P=<0.001; HPC: 13.7%, Z= 2.98, P=0.010, proportions *z*-test, Bonferroni-corrected). Responses to light onset were suppressed in multisensory trials compared to visual-only trials in a significant proportion of V2L cells (Fig. 3e; V2L suppressed: 10.9%, Z=2.58, P=0.020; V2L enhanced: 4.4%, Z=0.87, P=0.780; HPC suppressed: 21.1%, Z= 1.98, P=0.095; HPC enhanced: 10.5%, Z= 1.14, P=0.509, proportions *z*-test, Bonfferroni-corrected). Individual cells in S1BF and PER were not modulated by light onset (Fig. 3b-d; S1BF: 6.8%, Z=0.60, P=1.00; PER: 5.9%, Z=0.456, P=1.00), proportions *z*-test, Bonferroni-corrected).

**Figure 3:**
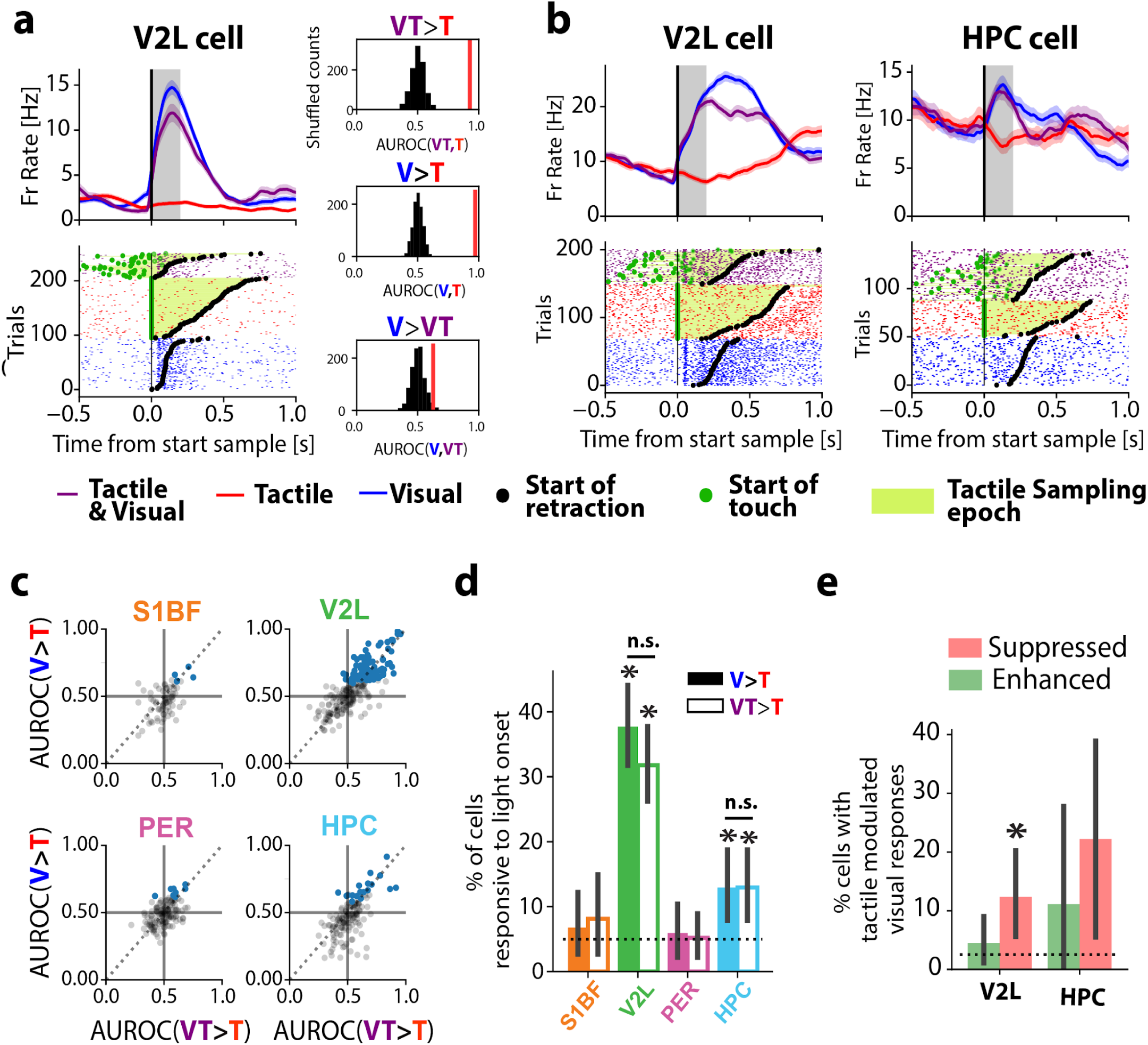
Visual responses of neurons recorded in all four areas during object sampling. **a-b)** Example cells modulated by light onset. Peristimulus time histograms (PSTHs) and spike rasters aligned on the start of object sampling are contrasting visual (blue), tactile trials (red) and multimodal (purple) trials. Light onset responses were quantified across the three different sampling conditions to account for potential confounds related to modality-specific sampling behavior. Multisensory trials are aligned to the light onset and the first touch in a trial is indicated by a green dot in the rasters. Retraction moments are marked by a black dot. This color coding holds in a, b, f and g. Vertical grey bar: time window used for analysis of light onset modulation. **a)** Right side: observed AUROC values (red) for firing rate differences for the example V2L cell and trial modality label shuffled chance level distributions (black). The values were measured by AUROC analysis on firing rates during the first 0.2 seconds after the light onset (grey shaded area). Cells with an AUROC value above 95% CIs of the shuffled distribution were considered to be significantly responsive to light onset. This example cell was modulated by light onset during tactile sampling (VT>T) as well as by light onset in absence of any whisker input (V>T). Tactile sampling suppressed light onset responses (V>VT). **b)** Two other example cells modulated by light onset. **c)** Overview of AUROC values for light onset responses of all cells per brain region. The majority of cells with significant AUROC values for light onset were observed in V2L, most often by responding to light onset in multisensory trials (x-axis, VT>T) as well as in visual trials (y-axis; V >T; blue dots indicate cells significant for at least one of the two modality contrasts). **d)** Percentage of cells per area which were modulated by light onset measured by AUROC analysis. Light onset was quantified by contrasting light onsets with combined touch to trials with touch in darkness (VT>T: empty bars; S1BF: 6.8%; V2L: 34.0% PER 5.9%; HPC:13.7%), or by contrasting light onsets in absence of touch to trials with only touch (V>T; filled bars; S1BF: 8.1%; V2L: 30.9%; PER 6.7%; HPC:13.7%). Black bars indicate 95% CIs. Asterisks indicate that the percentage of significant cells was higher than chance, (P<0.05, two-proportions z-test). This holds in e, h and j as well. **e)** Percentage of light onset responsive cells in V2L and HPC that showed tactile modulations of the light onset response (S1BF and PER are not shown due to lack of significant modulation, see d). For all cells that were significantly responsive to light onset (V>T), tactile modulation of the light onset response was quantified by contrasting visual responses with combined touch (VT) to visual responses in absence of touch (V2L Suppressed: 10.9%; V2L Enhanced: 4.4%; HPC Suppressed: 21.1%; HPC Enhanced: 10.5%).

### Tactile responses

Next, we quantified responses to individual whisker-object contacts by AUROC analysis on firing rates responses in the first 50 ms after individual touches. Cells in S1BF were significantly modulated by this early touch phase, whereas cells in other regions were not (Fig. 4d, e; S1BF: 22.2%, Z=3.93, P<0.001; V2L: 5.0%, Z=0.014, P = 1.00; PER: 5.3%, Z=0.179, P=1.00; HPC: 2.7%, Z=-1.904, P=1.00, Proportions *z*-test, Bonferroni-corrected). We additionally analyzed modulation of firing patterns by tactile sampling on a coarser time scale (from start of sampling up to retraction, defined by means of high-speed video tracking at 500 frames/s). On this coarse time scale a significant proportion of HPC neurons did respond to whisker touch (Fig. 4c; 20.1%, Z=4.30, P<0.001), proportions *z*-test, Bonferroni-corrected). We found no marked responses to whisker touch in S1BF, PER and V2L when tested on this coarse time scale (S1BF: 9.6%, Z=1.31, P= 0.380; V2L: 5.5%, Z=0.34, P = 1.00; PER: 3.0%, Z=-1.39, P=1.00). Subdividing HPC cells into the different subregions of origin (CA1, CA3, DG) did not reveal significantly different proportions of responsive cells between subregions for touch or light onsets (P=1.00, Proportions *z*-test, Bonferroni-corrected), therefore these HPC cells were pooled. The main result from these analyses is that tactile and visual responses evoked at stimulus onset – although expressed in S1BF (fig. 4e) and V2L (fig. 3d) – do not emerge in PER when rats sampled objects through vision and/or touch, despite the finding that such responses are, in fact, expressed in HPC (Fig. 3d, 4c).

**Figure 4:**
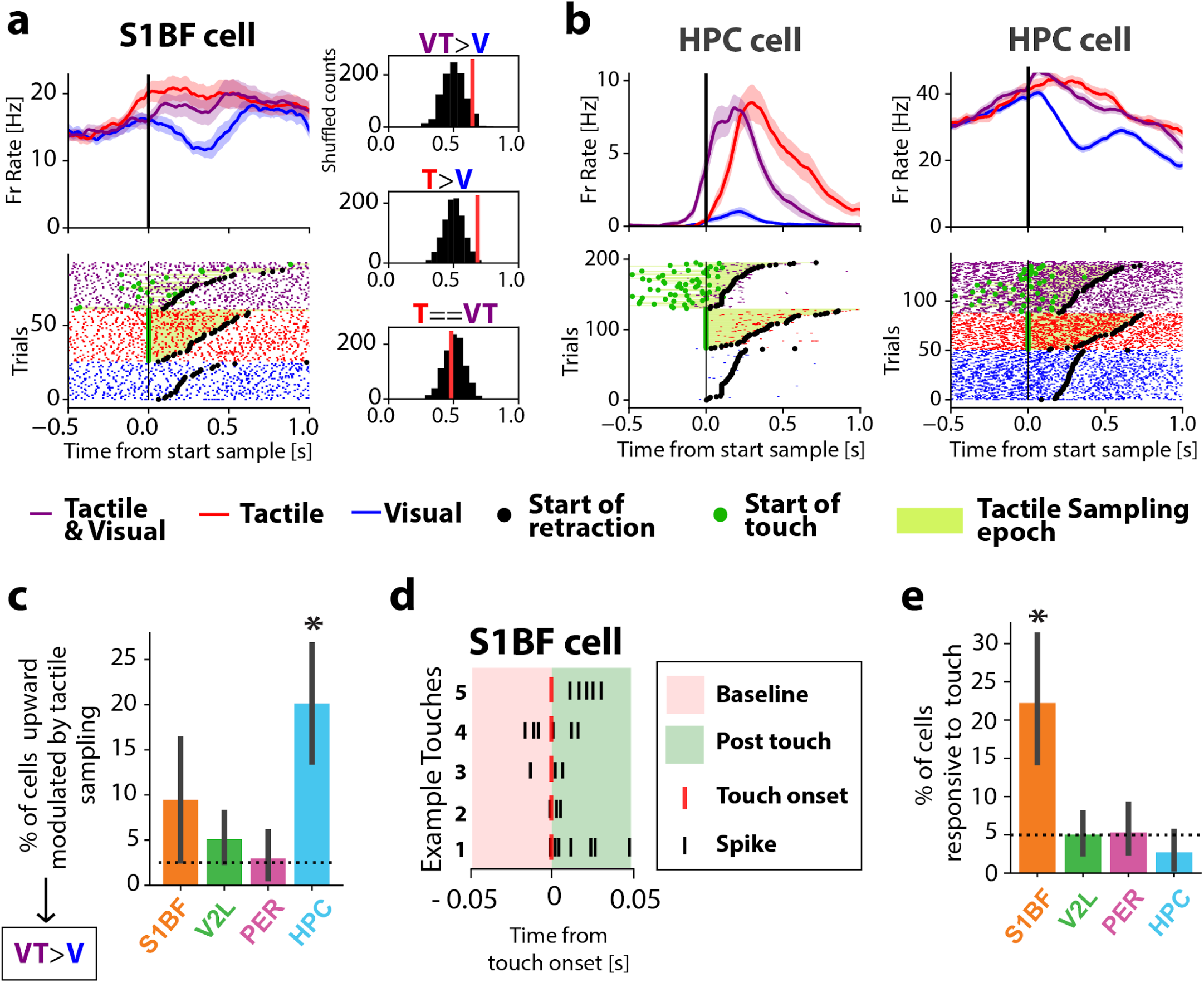
Tactile responses of neurons recorded in all four areas during object sampling. **a-b** Example cells modulated by tactile sampling. PSTHs and spike rasters contrasting visual (blue), tactile trials (red) and multimodal (purple) trials. Multisensory trials are aligned to the light onset. The first touch in a trial is indicated by a green dot in the rasters. Retraction moments are marked by a black dot. **a)** Right side: observed AUROC values (red) for firing rate differences for the example S1BF cell and trial modality label shuffled chance level distributions (black). The values were measured by AUROC analysis on firing rates during sampling, up to retraction from the object. Cells with an AUROC value above 95% CIs of the shuffled distribution of the VT vs. V comparison were considered significantly responsive to tactile sampling because these trials had identical illumination conditions. This example cell was modulated by tactile sampling regardless of the illumination (VT>V and T>V). Tactile responses were not modulated by the light onset (T==VT). **c)** Percentage of cells per area showing firing-rate enhancement by touch onset under identical illumination conditions, as measured by AUROC analysis on firing rates during sampling, up to retraction from the object (S1BF: 9.56%; V2L: 5.51% PER: 2.96%; HPC: 20.14%). Bars indicate 95% CIs. **d)** Spike times aligned to example touches (red) from one example S1BF cell. Touch onset responses were measured for each cell by contrasting the 50 ms following each touch onset (green background), with the 50 ms preceding the onset (red background) **e)** Percentage of cells per area which are modulated by touch onset in the first 50 ms after touch onset (S1BF 22.2%; V2L 5.0% PER 5.0%; HPC 2.5%). Bars indicate 95% CIs.

### Visual object representations in secondary visual cortex, but not in perirhinal cortex or hippocampus

The PER is sometimes considered a terminal area of the visual cortical stream, involved in object (or item) perception and recognition memory (Bartko *et al.,* 2007, Bussey and Saksida 2005, Norman & Eacott 2004; Dickerson and Eichenbaum 2010). To obtain reward during our task, the rats needed to sample objects and recognize the objects by vision and touch. We quantified how object identity was represented by individual cells in the hierarchy using a generalized linear model (GLM) that included choice side and object identity as predictors. The GLM quantified the influence of both predictors on neural responses over time, based on the instantaneous firing rates during all trials, aligned on sample start. We did this separately for tactile, visual and multisensory sampling, assuming that neural representations may differ depending on the sensory modality at hand. Cells were considered to be modulated by object identity if the GLM object coefficient magnitude was larger than what would be observed by chance, within 0.5 seconds after initiation of object sampling. This epoch was chosen because high object predictor magnitudes clustered during this epoch, and the choice side predictor became dominant thereafter (Fig. 5). V2L cells expressed significantly different responses depending on object identity during visual sampling, but not during tactile sampling (Fig. 5d,f; visual: 9.2%, Z=2.34, P=0.04) tactile: P =1.00; Proportions *z*-test, Bonferroni-corrected). In contrast, S1BF, PER and HPC cells were not encoding the object identity by firing rate during visual sampling behavior at all (Fig. 5e, f, S1BF: 5.1%, Z=0.05, P=1.00; PER: 3.6%, Z= -0.93, P=1.00; HPC: 3.5%, Z=-0.96, P=1.00). We did not find above-chance proportions of cells that encoded objects during tactile-only trials in any of the recorded regions (P>0.05 for all regions, Proportions *z*-test, Bonferroni-corrected). Additional control analysis confirmed that V2L cells with significant GLM object predictors tended to have higher AUROC values for object identity than for choice side (T(618) = -5.16, P<0.001; independent t-test). We verified that object correlates did not emerge later in time by applying the same analysis to a longer time window (0.5-2 seconds post-sampling). Thus, whereas V2L cells coded object identity, this coding was surprisingly absent in PER and HPC.

**Figure 5:**
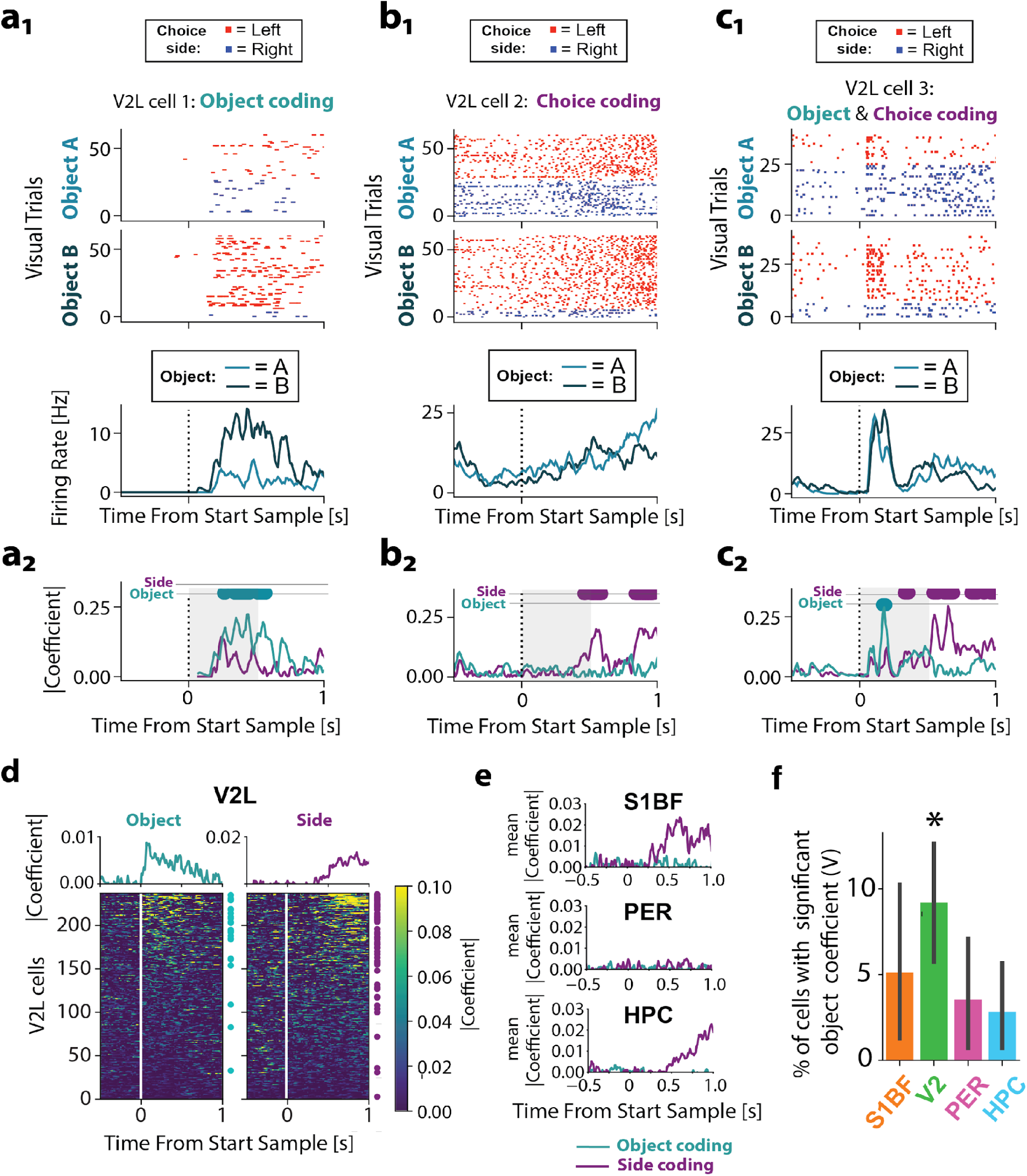
Visual object representations in V2L, but not in PER, S1BF or HPC: **a1-2)** Example V2L cell that is coding for object identity. This cell’s response to object B cannot be explained by choice behavior: note the raster differences between the two objects for only the left choice trials (for this rat the left choice was associated to object B). Top panels: Spike rasters of visual trials grouped by object identity, and color coded according to response side. Bottom panel: Mean firing rates over time for trials in which object A was presented versus trials presenting object B (visual trials only). Shaded bands indicate SEM. **b1-2)** Example V2L cell that is coding for choice side, but not for object identity. **c1-2:** Example V2L cell that first codes for object identity, and codes for choice side thereafter. **a2-b2-c2)** Absolute magnitudes of the GLM coefficients for the choice side and object identity for the same cells as in a1-c1. Colored dots above coefficients indicate significant encoding of the object identity (cyan) or choice side (purple; P<0.05 Benjamini-Hochberg corrected). The shaded grey areas indicate the time window used for subsequent calculation of object coding indices in **f**. **d)** Time course of object coding dominance and choice coding dominance in V2L. Heatmaps display the difference in predictor coefficient magnitude over time for all 237 V2L cells (Left: object, Right: choice). Colored dots on the right side of a heatmap indicate cells for which the object predictor (cyan) or choice-side predictor (purple) contributed significantly to the firing rate variance on any time point within the 0.5 seconds after starting to sample. The cells are ordered according to the maximal dominance of a given predictor (Left: object dominance, Right: choice dominance. **e)** Time course of the average difference in predictor coefficient magnitude from all S1BF, PER and HPC (compare to V2L, upper panels of (d)). **f**) Area-specific percentages of cells with significant GLM-based object modulation in visual trials (S1BF: 5.1%; V2L: 9.2%; PER: 3.6%; HPC:3.5%). Bars indicate 95% confidence intervals (CI). * indicates percentage of significant cells higher than chance (P<0.05, two-proportions z-test).

### Neural correlates of choice side in perirhinal cortex temporally align to reward site visits

In addition to the anatomical connections between PER and sensory cortical regions, PER also receives projections from subcortical motivation-related structures such as the substantia nigra, ventral tegmental area and amygdala, as well as from medial PFC (Burwell, Witter & Amaral, 1995; Kajiwara *et al*., 2003; Agster & Burwell, 2016). Previous studies demonstrated that PER cells carry spatial information during choice behavior in a task environment (Bos *et al*., 2017) and differentiate between choice sides most strongly at goal-specific locations (Ahn & Lee 2015). Corollary discharge and vestibular-proprioceptive inputs may be expected to arise upon initiation of locomotion towards reward goals, whereas representations of goal locations may arise upon arrival at the reward ports. We found that PER firing rates were not affected by choice side when rats initiated a turn towards the goal locations for reward when they retracted from the object (interval: 0 to 1 s from sample start; Fig. 6b). This absence of choice side coding contrasted to modulations observed in the HPC and the sensory regions, which did reflect changes in snout positions during opposed directions of self-motion when retracting from the object.

**Figure 6:**
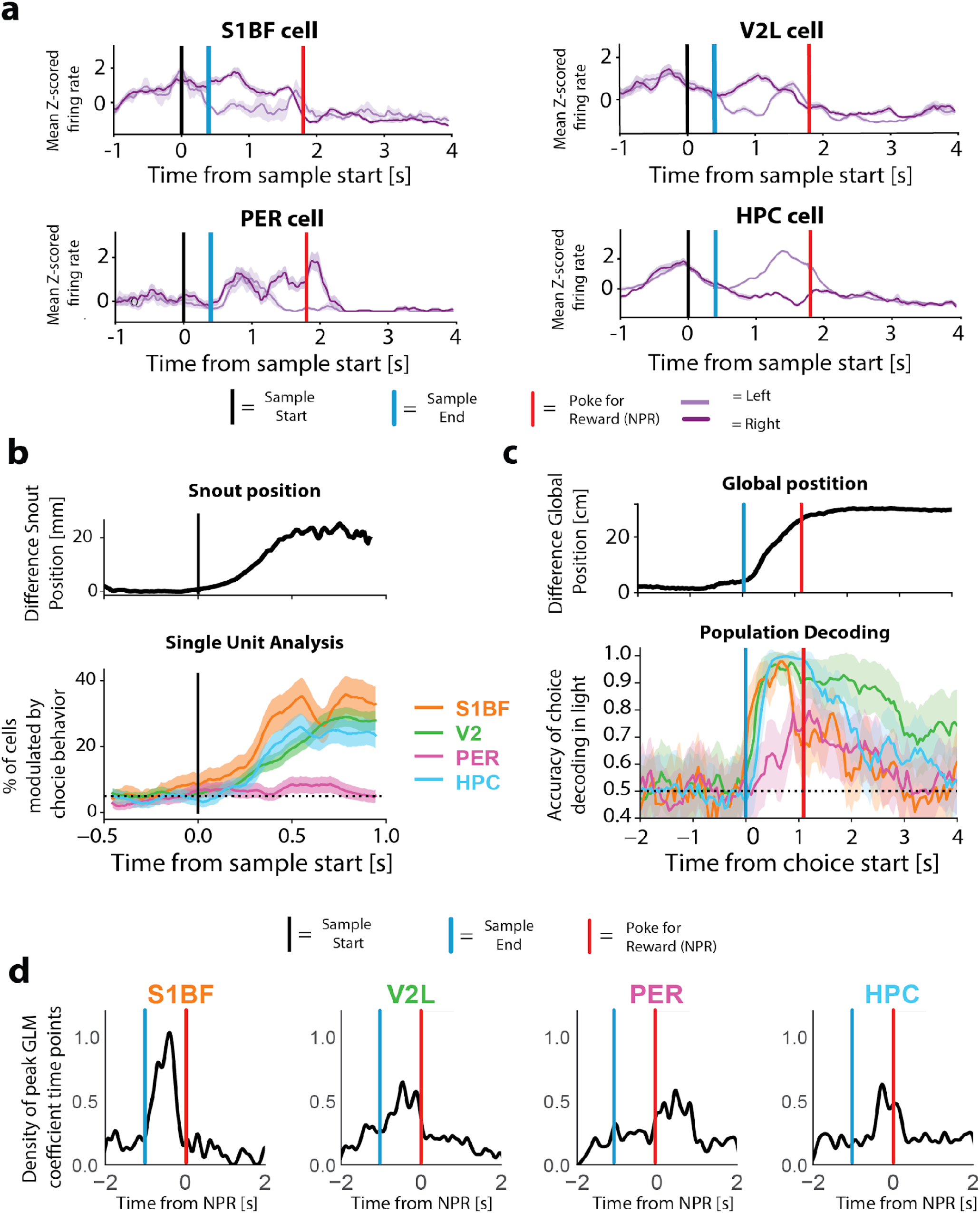
Temporal dynamics of choice side coding. S1BF, V2L and HPC cells were modulated mainly during the choice phase, whereas PER cells were mostly modulated during the outcome phase. **a)** Z-scored firing rates of example cells in the cortico-hippocampal hierarchy. Firing rates are linearly warped to align on the start of object sampling (black), the end of sampling, coinciding with the start of the left or right choice turn (blue), and the nose poke for reward (NPR; red) to account for unequal durations in object sampling and response latencies. **b)** Top: Average difference in snout position between left and right choice trials during object sampling behavior. Rats initiated a sideways motion towards the chosen side already within 1 second after sampling onset. Bottom: Percentage of cells significantly modulated by choice side, plotted as a function of time. Initial sideways motion towards the chosen side was represented prominently by cells in the sensory cortices and hippocampus, but not by cells in the PER. **c)** Top: Mean global position of two example rats on the elevated platform during choice behavior (black). Bottom: Temporal dynamics of population decoding accuracy of choice side, based on firing rates linearly warped for temporal alignment on the end of the sampling, marked by retraction from the object and the start of the left or right choice turn, and the poke for reward. **d)** Densities of poke-aligned times at which cells maximally differentiate between the choice sides, based on coefficients from a GLM that included choice side, object identity, sensory modality and trial outcome as predictors. Only cells with a significant choice side predictor are included in the densities.

Contrasting with the lack of specific coding during object sampling and retraction, we did in fact observe that PER cells preferentially discharge after the NPR (Fig 2c and 2e, S3; probabilities, S1BF: 0.176, Z=1.24 P= 0.426; V2L: 0.172, Z= 2.02, P= 0.086; PER: 0.281, Z=4.63, P < 0.001; HPC: 0.113, Z= -0.50, P= 1.00, tested against chance level of uniformly distributed events, proportion *z*-test, Bonferroni-corrected). We examined whether PER cells preferentially differentiated between the two goal locations for reward by quantifying the effect of choice side on firing rates over time at the population level. We constructed pseudo-populations that consisted of rate vectors from 100 neurons obtained in different sessions, for each area separately (see Methods). We then quantified how well a classifier performed in differentiating between the opposed choice sides based on the firing rates over time. The choice side of the animal could be accurately decoded from the population responses from S1BF, V2L and HPC as soon as the animals initiated their choice response, both in darkness and light. In PER, decoding accuracy of choice side was generally lower during locomotion, but peaked after the first NPR (Fig. 6c). Multiple task variables could potentially affect firing rates of individual cells during choice behavior. To disentangle the contribution of photic condition, choice (Left/Right), reward and object coding, we examined the relation between each of these task variables and neural responses with a GLM. Cells were considered to be selective for choice side if the GLM fit resulted in a significant choice side coefficient at any moment in the trial (see Methods). PER choice side coefficients preferentially peaked following the NPR, whereas cells in other regions differentiated between sides most prominently during the preceding choice phase, marked by locomotion towards the goal location (Fig. 6d). In total, 12.2% of PER cells encoded the choice side within one second after the NPR, as opposed to 8.0% in the second just before NPR entry. These results demonstrate that PER neural responses are more strongly correlated to the moments of (expected) reward delivery, compared to the HPC and sensory regions, which were modulated by choice side most strongly during locomotion.

### PER encodes choice and trial outcome when sampling for reward

Our results indicate that PER cells preferentially discharge after the poke for reward, and that the PER increasingly started to differentiate between choice sides when rats arrived at the goal locations for reward. The PER might integrate information on expected reward for a specific goal location with information on the actual outcome, reminiscent of prediction values and prediction errors in temporal difference reinforcement learning (TDRL) algorithms (Schultz *et al*., 1992; Sutton, 1988; Sutton and Barto 1998; Pennartz 1995). We scrutinized the period in which the rats experienced the actual trial outcome based on nose-poke and lick behavior. This ‘Initial Outcome Period’ (IOP) started when rats poked the reward port, after which they were confronted with the delivery of reward or the omission of it. The rats’ reactions were then measured by means of the first lick after the NPR, which we considered to be the moment that rats started to sample for the outcome. The IOP lasted until rats had started to sample for reward delivery by licking in 95% of all trials. Importantly, lick rate adjustments to the availability of reward (delivery vs. omission) occurred subsequently to this epoch (Fig. 7a). Thus, any differences in firing rate between rewarded and unrewarded trials after the IOP could not be differentiated from reward-associated motor activity (e.g. licks and reward port exits), whereas firing rate modulations during the IOP were unlikely driven by these confounds. Upon inspection of individual neurons, we noted that a subset of PER cells transiently increased their firing rates during the IOP depending on the actual trial outcome (Fig. 7b). In the other regions, cells generally fired according to the outcome only after the IOP and in a more sustained way (Fig. 7). We employed a GLM on the reward phase to disentangle the contributions of trial outcome, photic condition (object-focussed light off or on) and object identity to time-dependent modulations in firing-rate distributions for each individual neuron (Figs. 8a, b, c, S4). We quantified whether individual cells carried information about trial outcome earlier in PER than in other regions by comparing the summed reward coefficient magnitude over time between significant cells from the different regions. During the IOP, outcome coefficient magnitudes increased earliest in PER (Fig. 8d). PER cells had higher outcome magnitudes during the IOP compared to a pre-poke baseline (Fig. 8d; P=0.001; one-way Anova (F=8.385, P<0.001), with post-hoc Tukey test). Similarly, the number of cells that preferentially encoded the outcome during the IOP was higher than expected by chance only in the PER, whereas cells in other regions generally responded thereafter, during outcome-dependent behavioral adjustments (Fig. 8e, f; P< 0.05; proportion *z*-test, Bonferroni-corrected; S1BF: 7.9%, Z=-2.46, P=0.99; V2L: 10.9%, Z=-1.72, P=0.96; PER: 30.3%, Z=2.08, P=0.02; HPC:1.6%, Z=-10.52, P=1.00). During the IOP the firing rate of individual PER cells did not correlate to the delivery of reward itself, because the majority coded for the omission of reward (Fig. 8g; Z=3.285, P<0.001, proportion *z*-test: 18 error-up cells> 5 correct-up cells). These results indicate that PER cells mainly signal negative outcome, when rats started sampling for reward delivery.

**Figure 7:**
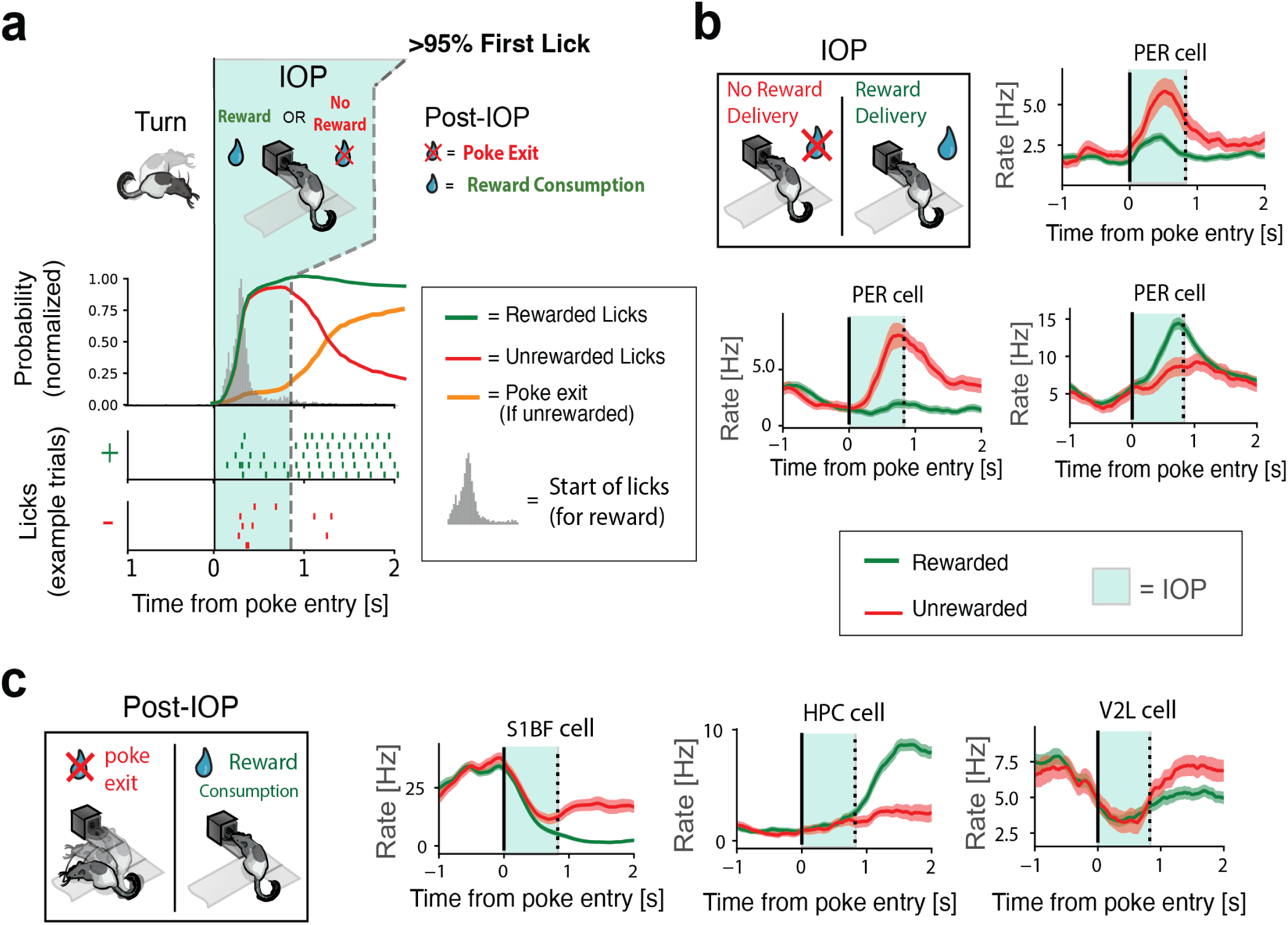
Trial outcome selectivity of perirhinal cells during reward delivery. **a)** Overview of poke and lick behavior during correct and incorrect trials. The Initial Outcome Period (IOP) was defined from the nose poke for reward up to the moment the rats had started licking in 95% of all trials. Even though a reward was only delivered in correct trials (and upon nose poke entry), rats initiated the lick response in all trials (rewarded and unrewarded) and adapted their licking behavior based on the presence of reward thereafter. Licking behavior thus typically consisted of initial licks to sample for reward delivery, followed by more licks for reward consumption in correct trials. **b)** Three example PER cells that differentiated between trial outcome during the IOP, when rats sustained their nose poke to sample for reward delivery. **c)** As opposed to PER cells, three example cells from S1BF, V2L and HPC are shown that became largely selective for trial outcome after the IOP, when lick and poke-exit behavior depended on the actual trial outcome (t=0 marks the onset of the IOP; vertical dashed line marks the end of the IOP).

**Figure 8:**
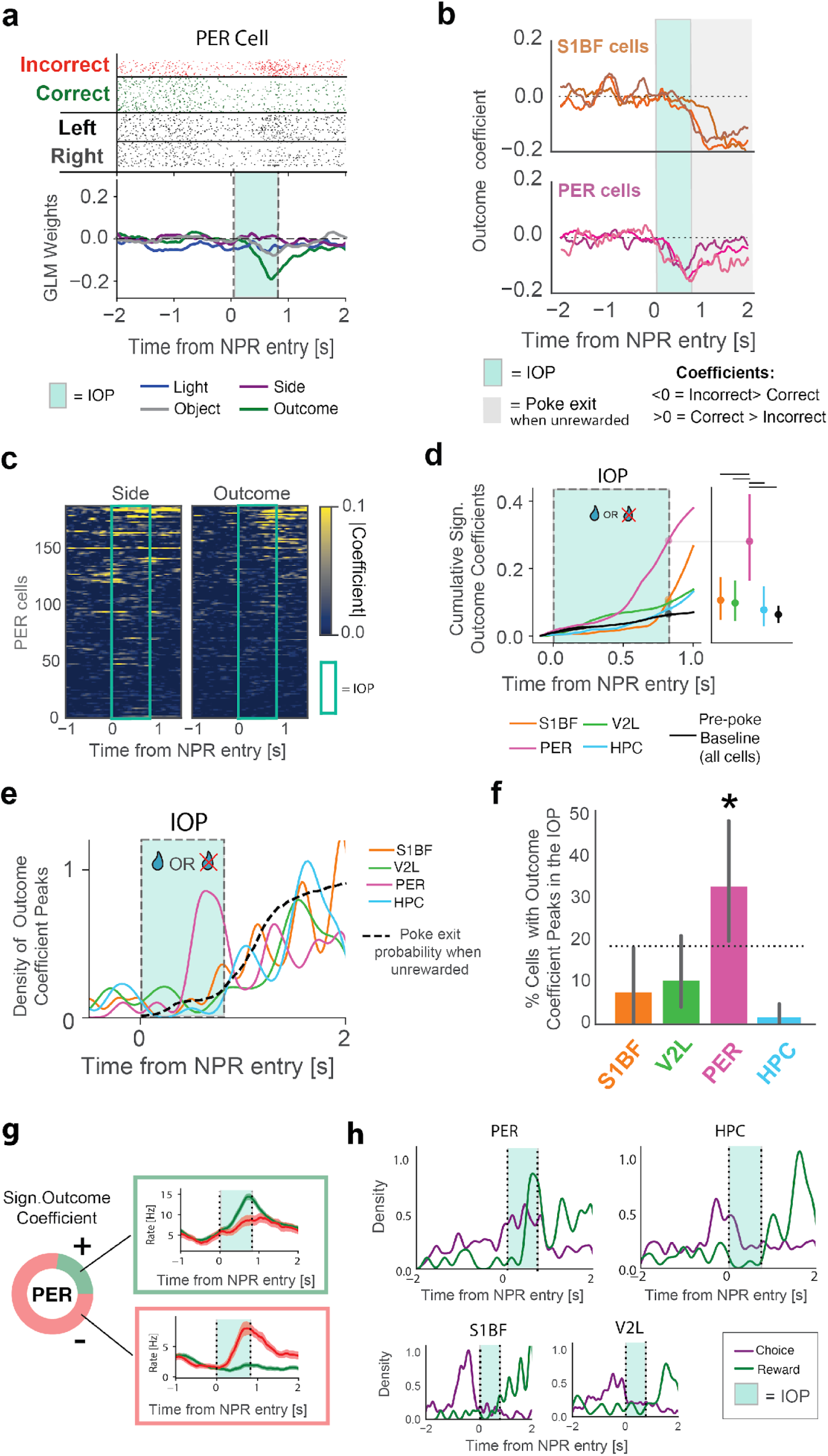
Simultaneous encoding of choice side and unexpected outcome in Perirhinal Cortex. **a)** One example PER cell revealing negative GLM weights for the outcome. The negative outcome weights reflect the increased firing rate during an unrewarded initial outcome period (IOP). Cyan area = IOP (as in b, d, e, g and h). **b)** Outcome coefficient time course of three example S1BF (orange) and PER (magenta) cells. Note that the outcome coefficients in S1BF cells increased after the IOP and are therefore likely driven by nose poke exits upon reward omissions. PER cells encoded the outcome earlier, when holding the nose poke for reward during the IOP (shaded cyan area). **c)** Time course of coefficient magnitudes for choice side (left) and outcome (right) for all PER cells. Cells are ordered according to their maximum outcome coefficient magnitude. Highest outcome coefficient magnitudes in PER clustered towards the end of the IOP (cyan outline), whereas the choice side was often encoded already before the nose poke for reward (NPR). **d)** Left: time course of cumulative outcome modulation following reward port entry. Outcome modulation is quantified by the cumulative sum of outcome coefficient magnitudes over time, averaged over all cells from a given area. Only cells with a significant coefficient for the outcome at any time bin are included. Right: cumulative sum of outcome coefficient magnitude at the end of the IOP. Colored vertical bars are CIs, horizontal bar for significant comparisons (P<0.05, One-way Anova (F=8.385, P<0.000) with post-hoc Tukey test). **e)** Time course of preferential trial outcome modulation for each area, based on the peak moment of the coefficient magnitude per significant cell. **f)** Percentage of significant cells that are modulated by outcome strongest during IOP as opposed to outside of the IOP (S1BF: 7.9%; V2L: 10.9%; PER: 30.3%; HPC:1.6%). Bars are permutation-based CIs, * for significantly higher percentages than uniformly distributed events (P<0.05, one-sided z-test of proportions). **g)** Left side: Proportions of significantly modulated PER cells having positive and negative outcome coefficients. Negative outcome coefficients resulted from higher firing rates during unrewarded IOPs compared to rewarded IOPs. Right side: Example cells responding to reward delivery or reward omission. Green and red traces indicate firing rates during rewarded and unrewarded trials respectively. **h)** Overlayed densities of poke-aligned times at which cells maximally differentiate between choice side (purple) and trial outcome (green), based on GLM coefficients. Only cells with significant coefficients for the given predictor (choice side or trial outcome) are included in the densities. Cells in the PER preferentially encoded both the choice side and the outcome (mainly reward omissions) when rats held the NPR during the IOP. The HPC and the sensory regions encoded outcome-related parameters after rats exited the reward site.

Because PER cells preferentially encoded the choice and outcome following the NPR, we compared the time course between choice side and trial outcome coefficients based on the GLM coefficients. Overlayed densities of preferential choice side and trial outcome encoding revealed that the PER simultaneously encoded the trial outcome and the choice side in the IOP, when sampling for reward delivery. Together, these results suggest that the PER integrates choice information with feedback on reward. This contrasted to the two sensory regions and HPC for which correlates were more tied to epochs of self-motion, likely arising from corollary discharge and sensory input during locomotion or from spatial coding (such as place fields for hippocampal cells).

## Discussion

In this study, we characterized the neural correlates of multisensory object sampling, choice side and trial outcome at prominent processing stages of the cortico-hippocampal hierarchy by conducting quadruple-area ensemble recordings in rats performing a multisensory object recognition task.

Our first hypothesis regarding the PER, pertaining to its potential contribution to object perception for familiar objects, predicted that object sampling would lead to sensory responses in S1BF, V2L and PER, and that the HPC would be less affected by the sensory modality at hand. Active touch evoked neural activity in S1BF and HPC, but not in V2L or PER. Light input at the start of visual sampling evoked neural responses in V2L and HPC, but not in S1BF or PER. Predominant responses to touch or light input were thus absent in PER, despite its known reciprocal anatomical connectivity to sensory cortical regions (Agster & Burwell, 2009; Burwell, Witter & Amaral, 1995). Importantly, PER was not systematically sampled along the full rostro-caudal axis. Tracing studies suggest that visual input mainly targets caudal PER, and tactile input primarily targets the rostral PER (Agster & Burwell, 2009). Modality-specific sensory input might thus be restricted to small subregions of the PER and could therefore be missed when recording cell samples. The majority of our PER recordings was performed in central-to caudal PER (Figs. 2a, S1), which has reciprocal projections with higher-order visual areas such as V2L but less so with S1BF (Agster & Burwell, 2009). Absence of visual responses in PER is therefore all the more striking, as is the contrast to HPC and sensory regions given that PER is often considered to be a relay station between sensory cortices and the rest of the MTL, including HPC (Naber *et al*., 1997; Doan *et al*., 2019; Fiorilli *et al*. 2021). Altogether, the present study supports the notion that sensory cortices are primarily modulated by their primary modality, and that multisensory (or amodal) processing is more abundant in the HPC compared to the PER and sensory regions considered here.

In visual trials, object identities could be decoded from V2L but not from PER, HPC or S1BF. These results show that visual representations of solid, 3D objects can exist without a robust object representation in caudal PER – even though this subregion receives many afferent fibers from higher visual areas (Agster & Burwell, 2009). Object identity was not prominently encoded by cells in any of the recorded regions during tactile sampling. Cells in S1BF may primarily represent low-level sensory features such as the angles and speed of whiskers upon object contact (Arabzadeh *et al*., 2004; Fassihi *et al*., 2020), much like cells in the primary visual system are mainly tuned by low-level visual features. Aside from modality-specific responses, object sampling did not lead to prominent amodal “object fields” in PER, as described in larger open field environments, where rats explored objects spontaneously (Burke, *et al*., 2012). As previously proposed, the behavioral task may require a spatial-navigational component for spatial firing fields to arise in PER (Ahn & Lee 2015; Bos *et al*., 2017; Fiorilli et al 2021). Notably, the objects we used were highly familiar to the rats and not explicitly configured to be ambiguous, which might be factors that lower the perceptual-mnemonic demands and recruitment of MTL structures such as PER (von Linstow Roloff *et al.,* 2016; Doron *et al.,* 2020). For instance, it is possible that sensory or invariant object representations only arise in PER when stimuli are novel signals, and this awaits future studies. Nonetheless, our results dispute an important perceptual-mnemonic role for PER in coding object-specific features when familiar objects are actively sampled and recognized.

### Perirhinal cortex and the neural coding of unexpected motivational outcome

The most prominent responses in PER were found after the animal’s arrival at the reward sites. First, PER was the only region in which cells preferentially encoded the choice side just after the NPR, as opposed to during the choice/locomotion phase (Fig. 5). Second, PER cells were responsive to the motivational outcome of a spatial choice during the reward delivery phase, just when rats started licking to sample for reward delivery. These signals were specific for the PER; correlates of trial outcome in HPC and the sensory neocortices were observed only when rats started to terminate the NPR (Figs. 6, 7d-e, S4c). In PER, signaling of reward omission was more often observed than when reward was delivered. This may relate to the larger degree of surprise associated with omissions than deliveries, because the range of correct responses ranged between 70-85% of trial types in which objects were presented (Fig. 1d). A systematic variation of (both negative and positive) surprise in trial outcome, applied against a background of variable reward history, must await future investigations.

Because PER firing responses peaked after reward port entry (when the subject was immobile), these PER responses (reported as population responses and GLM coefficients) are unlikely to be driven by locomotion, vestibular-proprioceptive inputs or by differential sensory flow accompanying body movements towards reward ports. Even though PER cells coded for choice side most strongly after the arrival at a goal location, the PER started to differentiate between sides (but not between outcomes) already before arrival at the goal locations (Figs. 6cd, 8h). This suggests that these cells anticipated the subject’s arrival at a given goal location: the closer to a given reward goal, the higher the firing rates for a specific population of PER cells. The relevance of reward delivery for PER firing is supported by earlier work demonstrating that PER cells lock onto large spatial segments of a maze associated with reward-and other task-related events; these firing fields generally peak at, or close to, goal locations (Bos *et al*., 2017). However, in this earlier study reward delivery was not accurately timed in relation to PER firing. Overrepresentation of motivationally relevant locations has been described in the HPC by the clustering of (usually small) place fields near reward sites (Hollup *et al.,* 2001; Gauthier and Tank *2018*; Lansink *et al*. 2012). The excess density of choice-side coding in HPC cells just before arrival at goal locations (Figs. 5e, S3b) aligns well with such overrepresentation. PER firing responses are notably distinct from place cells observed in HPC as they preferentially encode the choice side after the NPR, as opposed to place cells activated before or upon arrival at reward sites (Fig. 5d).

The observed neural correlates of outcome in PER are reminiscent of value and reward prediction signals in TDRL algorithms, neural correlates of which have been attributed to mesencephalic dopamine neurons (Mirenowicz and Schultz 1994; 1996; Schultz *et al*. 1997; Stalnaker *et al*., 2019; but see Rusu and Pennartz 2020). A single dopaminergic cell may encode both positive and negative prediction error, whereas PER cells in our study were found to code by way of reward-up or omission-up firing-rate responses. More recently, comparable reward value signals have been reported in the serotonin system, such as in the dorsal raphe nucleus which is well connected to the PER (Vertes, Fortin & Crane, 1999; Bromberg-Martin *et al*., 2010). Much like dopamine signals, sustained activity of dorsal raphe neurons can be driven by expected reward, and signal positive or negative reward prediction errors. In contrast to midbrain dopaminergic neurons which are phasically excited by positive errors and decrease firing activity upon negative errors, serotonergic neurons are primarily excited by punishment (Cohen *et al*., 2015). Encoding of punishment may be reminiscent of, but is different from signaling the omission of an expected reward, which is the predominant effect seen in PER neurons.

As compared to the study by Ahn and Lee (2015) on rat PER, we already noted some similarities in findings, but several points can be highlighted where the current study goes significantly beyond their results. First, we recorded from three other cortical areas in addition to PER, which allowed us to establish a positive control for object coding as compared to the null finding, viz. that PER, somewhat surprisingly, does not code object identity. Our finding that V2L neurons did code object identity (Fig. 5) shows that the solid, 3D objects and the task protocol we used were suitable for identifying object correlates in the neocortex, which was not shown by Ahn and Lee (2015), using 2D images as visual stimuli. Second, Ahn and Lee’s task deployed an auditory cue, delivered at the time of the choice response in their task, to signal whether the rat’s behavioral response was correct or not, which confounds the identification of choice correlates with both sensory input (tone) and tone-induced changes in reward expectation. Third, Ahn and Lee did not segregate the reward consumption phase from an earlier IOP period (as in Fig. 8), making it difficult to disentangle effects of reward from those of whole-body motor activity elicited by (non-)reward. These confounding factors may explain why these authors found relatively many PER cells to be modulated during the behavioral choice phase and less cells whose activity was related to reward or its expectation.

In line with Eradath et al. (2015), one of the primary objectives of our study was to characterize the neural representations of stimuli and their corresponding reward outcomes in PER. Eradath et al. (2015) found that cells in PER of macaque monkeys represent cue-outcome associations and temporal context. PER cells mainly represented the outcome type contingent on the cue and showed sustained activity from cue onset until the next trial. This representation depended on visual stimuli and was not present when rewards were given independently of cues. However, a noteworthy distinction in our study is the incorporation of multisensory stimuli, which provides a unique perspective on the encoding and representation of cue-outcome associations in real-world scenarios. Our results diverge from previous research by revealing that even with the inclusion of multisensory stimuli, clear object representations were not evident in the PER. This finding challenges the prevailing notion and suggests that other neural mechanisms or regions (i.e., V2L) may play a more prominent role in object percept formation.

The causal involvement of the rodent PER in object perception or memory has often been investigated by assessing impairments in detecting odd or novel items (Bartko *et al*., 2007; Albasser *et al*., 2010; Reid, Jacklin, & Winters, 2012). In the context of reinforcement learning, novelty is frequently viewed as a predictor of a possible reward, or as being rewarding in and of itself, which encourages exploration before actual appetitive benefits are realized (Akiti *et al*., 2022, Kakade and Dayan, 2002). The current study reports neural representations of rewarded locations (i.e., site where the NPR is produced) co-occurring with reward-outcome signals in PER, suggesting that PER lesions might disrupt the assignment of error-signals to unexpected items associated with novelty detection.

It may thus be hypothesized that PER facilitates learning and adaptive behavior by taking part in signaling unexpected reward events (be it as negative or positive surprise) for specific actions such as spatial choices, wherein in the interaction with serotonin and/or dopamine signaling needs further investigation. As elsewhere in the cortex, pyramidal cells of the PER use glutamate as neurotransmitter, and it is noteworthy that Reinforcement Learning based on deviations from expected reward can also be implemented in models using glutamatergic, Hebbian synapses (Pennartz 1997). The hypothesis that PER plays an important role in reward-dependent learning is supported by Doron *et al*. (2020) showing that PER outputs arriving in layer 1 of rodent somatosensory cortex are critical for learning associations between stimuli and reward and that these become unnecessary for correct task performance once the task rule has been acquired.

Altogether, the present study challenges the notion PER is involved in perceptual processing, suggesting that deficits in visually guided behavior observed after PER lesions may be primarily due to deficits in predicted reward outcomes and novelty detection. This interpretation is supported by the absence of such reward-based feedback in V2L (as well as the other two areas under scrutiny). It further suggests that the final stage of visual object perception takes place upstream of PER (e.g. V2L or TeA), while the PER plays a critical role in learning and remembering associative relations among events but does not contribute significantly to object perception. Further studies examining learning under parametric variations of reward parameters will be required to determine how motivational feedback and error signaling in the PER compare to adjacent structures in the MTL, such as the postrhinal and entorhinal cortices. More work along these lines is needed to characterize the input-output wiring of reward-related cells in PER in more detail, and to quantify how signaling of unexpected outcomes and errors evolves over the course of learning.

## ACKNOWLEDGEMENTS

The authors would like to thank Jeroen Bos and Laura Mourik Donga for their technical support for surgeries and hyperdrive assembly. We additionally thank Charlotte Oomens and Carien Lansink for their input and advice on the behavioral paradigm and training procedure. We are grateful for the support provided by the Technology Center of the University of Amsterdam, in particular by Sven Koot, Tristan van Klingeren, Gerrit Hardeman, Tjeerd Weijers, Udo van Hes and Alix Wattjes. We thank C. Rossant, members of the Cortex Lab (UCL) and contributors for Klusta and Phy spike sorting software and the HBP data services for data curation. This project has received funding from the European Union’s Horizon 2020 Framework Programme for Research and Innovation under the Specific Grant Agreement No. 945539 (Human Brain Project SGA3) to C.M.A.P.

## CONTRIBUTIONS

J. F. and C.M.A.P. designed and planned the experiments; J.F., M.D.Q. and R.B. trained the rats on the task; J.F. and G.H. did the surgeries; J.F. performed the quad-drive tetrode recordings, with support from M.D.Q and R.B.; T.R. programmed preprocessing pipelines and curated all data; I.R. and J.B. mapped and integrated the position of recording sites in a common reference atlas; J.F. and P.M. analyzed the data with support from C.M.A.P.; C.M.A.P. and J.F, wrote the paper in collaboration with the other co-authors; C.M.A.P. supervised and coordinated the project.

## Methods

### Subjects

Data was collected from four 28–44 week old male Lister Hooded rats (obtained from Envigo, The Netherlands). Rats were socially housed under a reversed day/night cycle (lights off: 8:00 a.m., lights on: 8:00 p.m.) and food restricted to maintain their body weight at 85% of the ad libitum growth curves of Harlan, taking Rolls and Rowe (1979) as a reference. All experiments were performed in accordance with the National Guidelines on Animal Experiments and were approved by the Animal Experimentation Committee of the University of Amsterdam.

### Apparatus

Behavioral training was performed in a darkened room on an elevated T-shaped platform (30W x 35L x H60 cm, 60 cm from floor) with a reward well for sucrose solution (15%) delivery at opposed sides (30 cm distance; Fig.1a). The training apparatus was fully automated with custom-written MATLAB code on a Windows PC, and hardware was controlled through a Field Programmable Gate Array (FPGA) and an Arduino. A pneumatic door prevented access to the objects during the intertrial interval (ITI). Nose pokes into reward ports (4W x 4H cm) and lick events were detected by two different IR phototransistors in each reward port (Fig. S2e). Reward was delivered by a syringe pump (Razel, USA). The on-and offset of the object sampling epoch were estimated during the task by phototransistors in front of the object, but more precisely defined for analysis afterwards by means of video tracking. The two reward ports were located on either side of the sampling platform, facing each other at a distance of 30 cm. To analyze object-sampling behavior, an infrared high-speed camera was mounted 40 cm above the rat’s head (500 frames/s; M3 Camera from Integrated Design Tools, USA, with 50 mm/F0.95 lens, Navitar, USA; Using MotionStudio software, Integrated Design Tools, USA). Object sampling, detected with phototransistors and infrared beams, triggered the high-speed camera to write 900 video frames for 1.8 secs to the PC. Background illumination of diffuse IR light was reflected from 5 LED arrays (Sygonix, 310 mA, wavelength: 850 nm) to the camera by a mirror located underneath the sampling region. White semi-transparent Perspex acrylic was used to diffuse the IR light coming from the side before being reflected to the camera. A second infrared camera (SONY EVI-D100P), placed 150 cm above the platform, allowed the experimenter to monitor the animal’s overall behavior on the elevated platform. A single IR light (Sygonix, 310 mA, wavelength: 850nm) mounted 40 cm above the platform weakly illuminated the maze for recording images on this camera.

The 3-dimensional objects consisted of plastic LEGO configurations which were mounted on black H11 x W20 cm rectangular acrylic background sheets with hot melt adhesive. Objects were randomly presented using a rotating stepper motor at the start of each ITI. The objects were randomly rotated before bringing the objects into place for an upcoming trial, in order to prevent that sounds related to stepper motor rotations would enable the animal to predict the upcoming object and associated reward location. In visual and multisensory trials, an object-focused white LED light was triggered upon object sampling. In tactile and multisensory trials, the objects were presented at a gap distance of 15-16 cm away from the elevated platform, which limited object contact to whiskers only (at this distance the object was typically at a distance of 0.5-3.5 cm from the snout during tactile sampling, see also Harris *et al*., 1999). In visual trials and catch trials, objects were presented further away (22-23 cm from platform), which made it impossible for the rat to reach the object with its whiskers.

Tactile object sampling was thus controlled by the gap distance to the object, and visual sampling was controlled by light illumination in an otherwise dark room. Computers, acquisition systems and experimenter were situated in a separate room from the darkened experimental room to prevent light leakage.

### Behavioral Procedure

We recorded four male Lister Hooded rats that were trained on a two-alternative forced-choice (2-AFC) object discrimination task. To correctly perform a trial, rats had to sample one of two possible objects, retract from the object-sampling port and subsequently make a choice by nose-poking into one of two reward ports, placed on the left and right side of the object-sampling port (Fig.1). The object-side associations were counterbalanced over the four rats, but the objects remained unaltered. A trial started after an intertrial interval (ITI) of 12 seconds, after which a pneumatic door opened (Fig.1a) and access to the object became possible. The modality (visual, tactile or multisensory) was selected randomly per trial, but was repeated after an incorrect response to discourage any preferred modality bias during a session. Rats were implanted and recorded when they reached a stable performance of at least 70% correct for each trial type during seven consecutive days.

### Surgical procedures and electrophysiology

Recording areas were targeted in the right hemisphere with a custom-built microdrive that contained 36 individually movable tetrodes (Nichrome wire, California Fine Wire, diameter: 13 μm, gold-plated to an impedance 300– 800 kΩ at 1 kHz), four of which were used as reference (Lansink *et al*., 2007; Bos *et al*. 2017). For two out of four rats, tetrodes were collected in four bundles, each of which targeted towards PER, dorsal HPC (CA1, CA3 and DG), V2 and S1BF (PER area 35/36; coordinates: -6.0 AP, 6.5 DV, 7.0 ML, dorsal CA1 and CA3 coordinates: -3.48 AP, 2.0 ML, 2.5 DV, V2L coordinates: -6.0 AP, 2.8 DV, 5.8 ML, and S1BF coordinates: -3.0 AP, 5.0 ML, 2.8 DV). For the other two rats, the hippocampus bundle was omitted; tetrodes from this bundle were appended to the V2L and PER bundles. Rats received pre-operative, subcutaneous injections of the analgesics buprenorphine (Buprecare, 0.04 mg kg−1) and meloxicam (Metacam, 2 mg kg−1) as well as the antibiotic enrofloxacin (Baytril, 5 mg kg−1). Anaesthesia was induced with 3.0% isoflurane, and was kept on 1.0–2.0% isoflurane during surgery. A heating pad was used to maintain body temperature. Lidocaine was applied directly on the periosteum for additional, local analgesia before exposing the skull. The skull surface was thoroughly cleaned with 3% hydrogen peroxide solution, and washed thrice with saline. Six screws, from which one served as ground (occipital bone), were inserted in the skull to improve the stability of the implant. After craniotomy and durotomy were performed, the drive was positioned using a stereotaxic holder. The craniotomy was then sealed using silicone adhesive (Kwik-Sil), and dental cement (Kerr-total adhesive Opti-bond Solo Plus and 3M Unitek Transbond) was applied to anchor the drive on the skull and screws (Bos *et al*., 2017; Lansink *et al*., 2007). Post-operative care included a subcutaneous injection of meloxicam on the two days following surgery, and Baytril one day after surgery. From the third day after surgery, tetrodes were gradually lowered towards their target regions on a daily basis. Recording locations were estimated based on the number of turns to the tetrode guiding screws, and by assessing the Local Field Potential (LFP), sharp wave ripples in CA1 and CA3, and spike signals during signal acquisition.

Neurophysiological signals were continuously acquired at a sampling rate of 32 kHz with a Digital Lynx SX 144 channel system (Bozeman, MT) and high pass filtered at 0.1 Hz. The signals were pre-amplified with a headstage before being fed through an automated commutator (Neuralynx). Spike detection and sorting were performed offline using Klusta and manually curated using Phy GUI (Rossant *et al*., 2016, also for optimized spike detection and clustering parameters). Clusters were included as units based on their spike waveforms, autocorrelation and stability over time. A unit was considered a single-unit if less then 0.5% of its spikes occurred in the refractory period (1.2 ms) and its isolation distance to other spike clusters was above 5. Only cells with an average firing rate higher than 0.5 Hz during trials (8 seconds pre-to 5 seconds post poke for reward) were included in single-unit analysis. Units that passed the above-mentioned criteria, but no other cases, were considered single units in the analysis. Spiking activity was aligned to events of interest and then convolved using an Alpha kernel (sigma=100 ms, step=10 ms), unless specified differently.

### Histology and anatomic registration of tetrode locations

On the final recording day, electrolytic lesions were made at the tip of each tetrode by passing current (18 uA for 2 s) through two leads of each tetrode. Approximately 24 hours after lesioning, animals were deeply anaesthetized with Nembutal (sodium pentobarbital, 60 mg ml^−1^, 1.0 ml intraperitoneal; Ceva Sante Animale, Maassluis, the Netherlands) and transcardially perfused with a 0.9% NaCl solution, followed by a 4% paraformaldehyde solution (pH 7.4, phosphate-buffered). After post-fixation, transversal sections of 40-50 μm were cut using a vibratome or sliding microtome and stained with Cresyl Violet to reconstruct tetrode tracks and localize their end-points. To allow direct comparison of recording regions between rats, we extracted spatial coordinates for all tetrode locations after registering the histological section images to the Waxholm Space Atlas of the Sprague Dawley rat Brain v3 (Research Resource Identifier (RRID): SCR_017124; Papp *et al*., 2014; Bjerke *et al*., 2018) using the QuickNII software (RRID: SCR_016854; Puchades *et al*., 2019).

### Video-tracking

We tracked whisker touches, snout and global position of the whisker array at either side of the snout, by training DeepLabCut (DLC; Mathis *et al*., 2018) on 548 video frames originating from 15 different sessions (N=4 rats). Video frames for manual annotations of head and whisker positions were picked from different sessions by k-means clustering based on frame appearances (Mathis *et al*., 2018). This ensured that labeling was performed on frames that looked different. We additionally hand-picked 260 frames from sessions that varied in snout position, sharpness, illumination and magnification. For quantification of whisking behavior, we tracked the rostral, caudal and midline of the whisker array at either side of the snout. We verified the tracking accuracy by (i) computing the median distance between tracked point and real annotated point (Mathis *et al*., 2018); (ii) plotting the whisker angles, head angles and moments of touch, and verifying that these followed the expected rhythmicity and spatial dimensions (rats whisk/palpate at a frequency of approximately 8 to 20 Hz, (Berg and Kleinfeld, 2003; Mitchinson *et al*., 2007; Sachdev *et al*., 2001); (iii) and visually inspecting videos with overlaid tracked whisker angles, tracked snout, and tracked touches (S2a, b, c, SI Video). Because the timing of individual whisk-touches was especially important to us, we additionally manually labeled all individual video frames from 5 sessions as having a whisker-touch in it or not. Comparisons between DLC-based whisk-touches and manual whisk touches revealed a high degree of overlap (Fig. S2d).

### Lick-rate analysis

To quantify the behavioral changes occurring in response to reward delivery, we computed a separate lick-PETH for rewarded and unrewarded nose-pokes into the reward port (Fig. 7a). Based on this licking behavior, we identified an epoch in which rats had started to sample for the reward in 95% of the trials (measured by means of the first lick after nose-poke entry. We identified this epoch as the ‘Initial Outcome Period’ (IOP), because rats started to exit the nose-poke site upon omitted reward delivery only after this point. Thus, significant outcome coefficients later than this epoch could also be explained by behavioral changes in lick patterns and poke behavior following reward sampling, whereas modulations within this epoch could not.

### Light-and touch onset modulation indices

We initially quantified responses to light- and touch onsets by comparing average firing rates during object sampling between visual, tactile and multisensory trials. Here, we restricted the analysis to a response window of 200 ms after the start of object sampling, thereby including the light and tactile onsets and limiting influences related to retraction movements. For every neuron, we used the area under the receiver operating characteristic curve (AUROC) to measure if sensory onsets from a given modality evoked higher firing rates compared to when this input was absent (e.g. for light onset responses we asked if VT>T and if V>T). Statistical significance of sensory differentiation was assessed by 1000 permutations of the trial modality labels to obtain a distribution of surrogate indices. AUROC values were considered to be significant for the tested modality when they fell above the one-sided 95% confidence interval of the surrogate distribution and the corresponding neurons were then considered to be modulated by the tested modality (see Fig. 3a for an example of visual modulation, and Fig. 4a for tactile modulations). For all cells that were significantly responsive to light onset we additionally quantified tactile modulations of the light onset responses by contrasting visual responses with combined touch (VT) to visual responses in absence of touch (V). A cell was considered to be suppressed or enhanced when its AUROC values fell respectively below or above the two-sided 95% confidence interval of the surrogate distribution.

To quantify very transient touch onset modulations, we identified clearly defined whisker touches of relatively long durations (of both left and right whiskers combined). This was achieved by selecting touch onsets for which at least 60% of the frames in the subsequent 50 ms contained a touch. Additionally, touch onsets were required to have a 50 ms pre-touch period with at most 30% of video frames displaying a touch. We selected sessions in which this procedure identified at least 30 touch onsets. For every neuron, we next binned spikes in the 50 ms before each touch onset (pre-touch) and after each touch onset (post-touch, Fig. 4d). The touch-onset modulation (TOM) index consisted of the AUROC values of pre-touch versus post-touch spikes. We then randomly permuted the pre-touch and post-touch labels 100 times per touch, and recomputed the AUROC values to obtain the null distribution. TOM index values were considered significant when they fell outside the 95% confidence of the surrogate distribution.

### Object coding

For every neuron, firing rates were aligned to the start of sampling and convolved as described earlier (see: Surgical procedures and electrophysiology). A GLM (see below) was used to disentangle the contribution of choice side and object identity, separately for visual and tactile trials. We restricted the significance estimation to the first 0.5 seconds after the start of sampling as we assumed that correlates of object identity would emerge during, or shortly after object sampling and recognition measured by means of whisker and snout tracking (for the object encoding over time, see Fig. 5d). We additionally verified that cells with significant object predictors also tended to have higher AUROC values for the object identity compared to the choice side, computed over the average firing rates from our analysis time window (see Results).

### Pseudopopulation decoding of choice side and reward outcome

Because multi-units can contribute to a population code, we included all manually curated units including those that did not meet the criteria for single-units in the population decoding analysis (N units: S1BF=203, V2L=488, PER=257, HPC=307). The pseudopopulation decoding of left and right choice was performed by randomly drawing 10 left tactile or visual and 10 right tactile or visual trials out of each recording session (recording sessions with fewer than 10 trials per type were excluded), and randomly selecting 100 cells across all sessions available from all subjects. For selected cells and selected trials, spikes were binned in 300 ms time bins, advanced with increments of 50 ms, and subsequently linearly warped to align retraction and reward-site pokes to the median event times computed across all sessions (median times were computed separately for visual and tactile trials). Then, at each time bin, we decoded left and right choices from spiking activity of the selected cells with a random forest classifier with 200 trees, using a 3 x 3 cross-validation routine, and computed the accuracy score of the predictions (Bos *et al*., 2019; Glaser *et al*., 2020). This entire procedure was repeated 500 times, each time randomly drawing the trials and cells. Reward outcome decoding (correct versus incorrect trials) was performed similarly, with a few minor differences. We used 20 trials of each class, and finer increments of the time window of 25 ms. Spike times were not warped but only aligned to reward-site pokes across trials.

### Generalized Linear Model

During object sampling or reward consumption, multiple variables may affect firing rates of individual cells. To disentangle the contribution of photic condition, choice (Left/Right), reward and object identity, we examined the linear relation between each of these task variables and the neural responses with a Generalized Linear Model (GLM; *Statsmodels* toolbox). We used a Poisson regression to approximate firing rate statistics (Fig. S4b). Single-unit activity was aligned to the event of interest (Start sample or Nose Poke for Reward, NPR) and convolved as described earlier. The GLM was fitted to describe the relationship between the binary trial variables and single-unit firing rate distributions independently on each time point. Accuracy of the GLM model was scored by predicting the firing rates of each cell for all trials on each time point, based on the coefficients through cross-validation, and by calculating the proportion of explained variance (R^2^ score) between predicted and actual firing rates (1.0 being the highest possible score). We verified the accuracy of the model fit by comparing R^2^ scores of time bins that had at least one significant predictor to time bins without any significant predictor contribution, as well as to time bins from a baseline epoch (−8 to -3 s before the poke), or by comparison to chance-level accuracies obtained by randomly permuting trial labels independently at each time-bin 500 times (Fig. S4). Statistical significance of a predictor at a given time point was determined with Z-statistics by comparing the explained variance against a model fit in which the tested predictor had no relation to the firing rates. P-values over time for a given cell were corrected by the Benjamini-Hochberg Procedure.

### Statistical significance of proportions of neurons

Statistical significance of cell proportions was determined according to z-test for proportions:

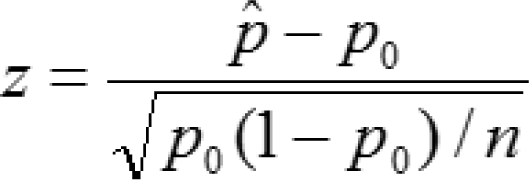

Where ^p^^ is the observed proportion of significant cells during an epoch of interest, and p_0_ the proportion of significant cells during chance level distributions, n is the total number of cells.

## Figures

## Supplementary Figures

### Supplementary Videos 1 and 2

**Figure S1.**
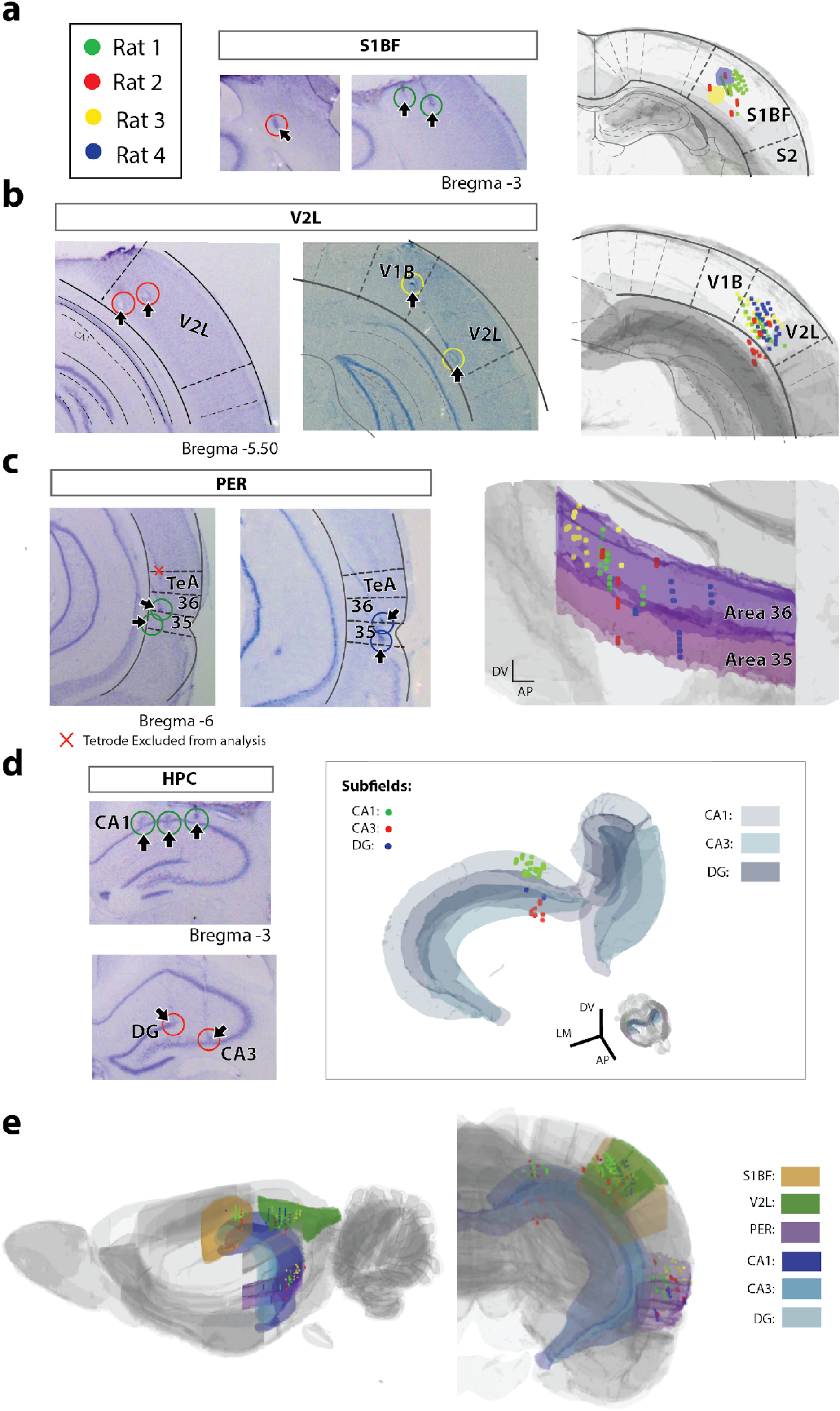
Histological verification of recording locations: **a-b-c-d-e)** Examples of histological sections and their locations in the three-dimensional Waxholm space reference atlas. Recording sites (dots) and tetrode endpoints (circles) are color-coded by rat and displayed separately for each target region. Arrows and circles indicate the tetrode endpoints in the histological sections **a)** Somatosensory Barrel Cortex (S1BF). **b)** Lateral Secondary Visual Area (V2L). **c)** Perirhinal Cortex (PER). Red cross indicates an endpoint of a tetrode that was excluded from analysis because histological verification revealed it ended up in temporal association cortex (TeA) instead of PER. **d)** Hippocampus (HPC) color-coded for subfields: CA1, CA3 and DG. **e)** Medial (left side) and coronal (right side) overview of the three-dimensional mapping of recorded locations on the Waxholm space reference atlas, for the four different target regions.

**S2.**
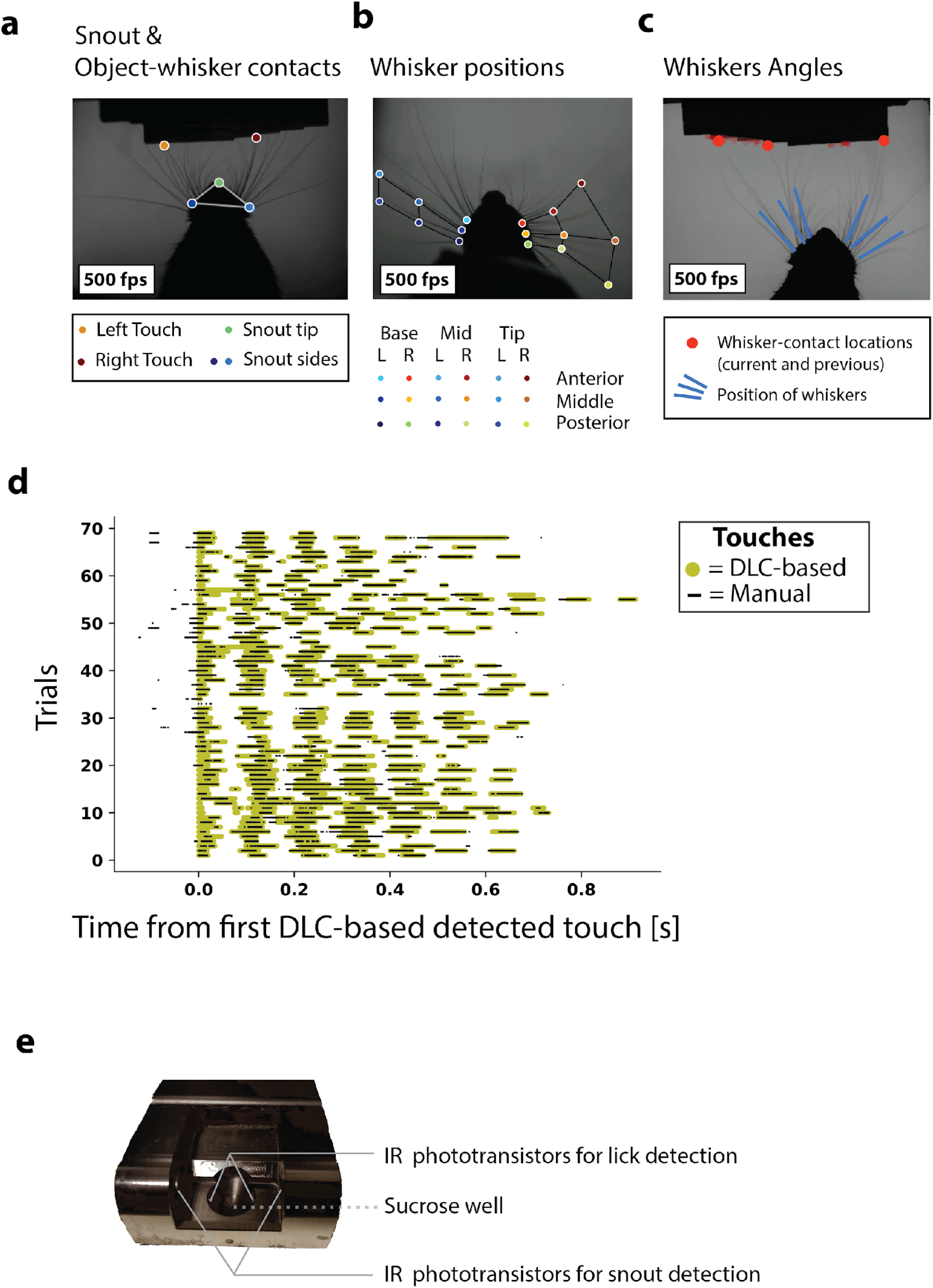
Video-tracking examples of whisking behavior and picture of reward consumption sensors. **a)** Example frame with overlayed position of snout and whisker-object contact detection for the left and right whiskers, tracked and displayed with DeepLabCut (DLC) software. Contact moments were used to determine the onset of tactile object sampling. fps: frames per second. **b)** Example frame with tracked positions of whisker base, mid and tips (proximal to distal) for the anterior, middle and posterior parts of the whisker arrays. Collectively, these positions allowed us to track global positions and angles from the two whisker arrays. **c)** Video frame with overlayed global position of whisker arrays (blue) and detected object-whisker contact points during the trial (red). **d)** Comparison between automatic (yellow; DLC) and manual (black) detection of whisker-object contacts during an example session. Moments of touch are aligned to the first touch detected by DLC in a given trial (plotted at time 0). **e)** Picture of a sucrose solution delivery well and the infrared (IR) phototransistors for nose poke and lick detection.

**S3:**
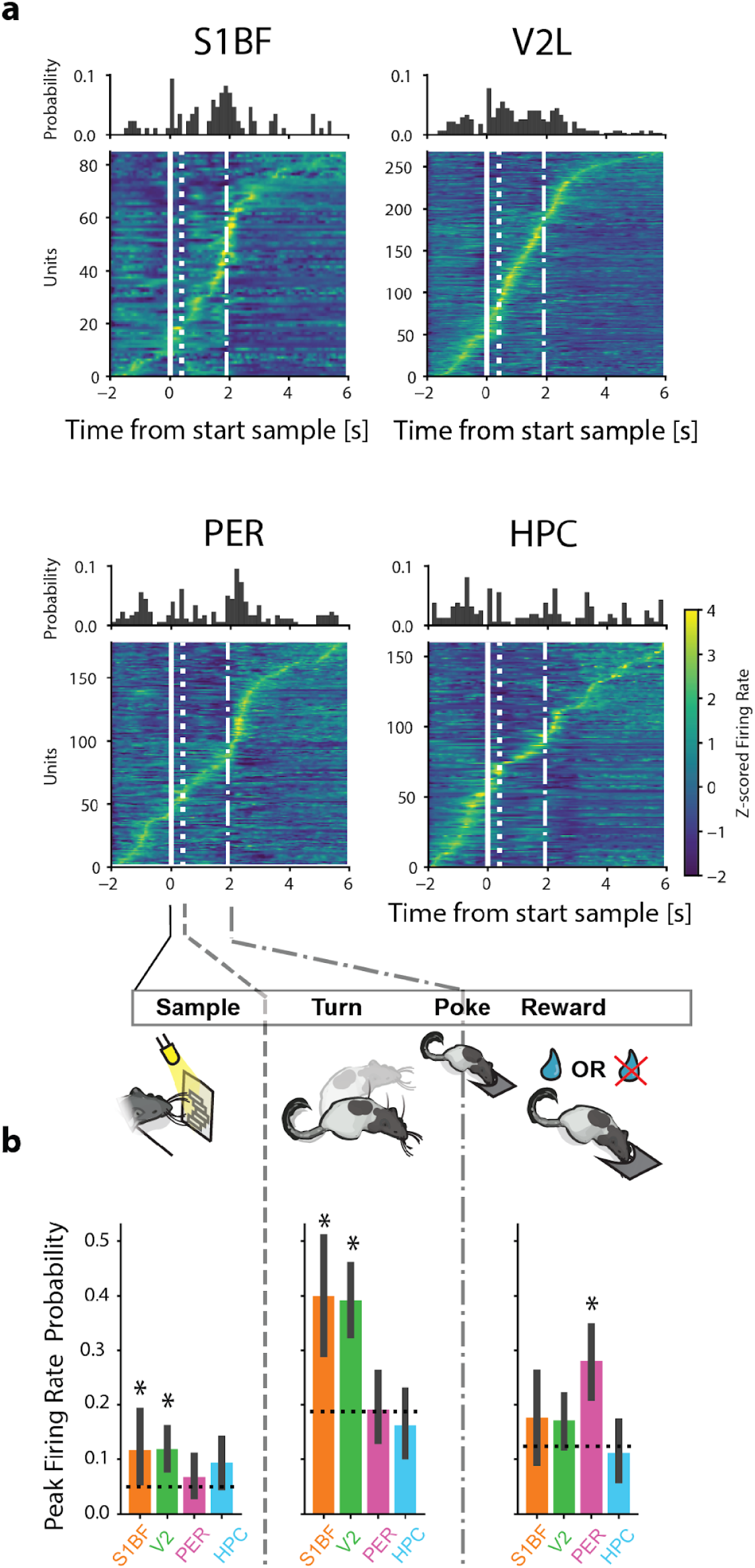
Perirhinal neurons preferentially discharge during the reward delivery phase as opposed to object sampling. **a)** Overview of task-phase modulation per area. Histograms indicate the distribution of probabilities in which units discharged maximally. Heat-maps are linearly time-warped Z-scored firing rates of cells over time (all the recorded single-units included). Cells are ordered by the moment of peak firing rate relative to sample onset. **b)** Probability of peak firing rates per task epoch. Bars are 95% CIs, dashed lines indicate chance levels given homogeneously distributed events over time. Asterisks indicate significant above-chance probabilities (P<0.05 two-proportions z-test).

**S4:**
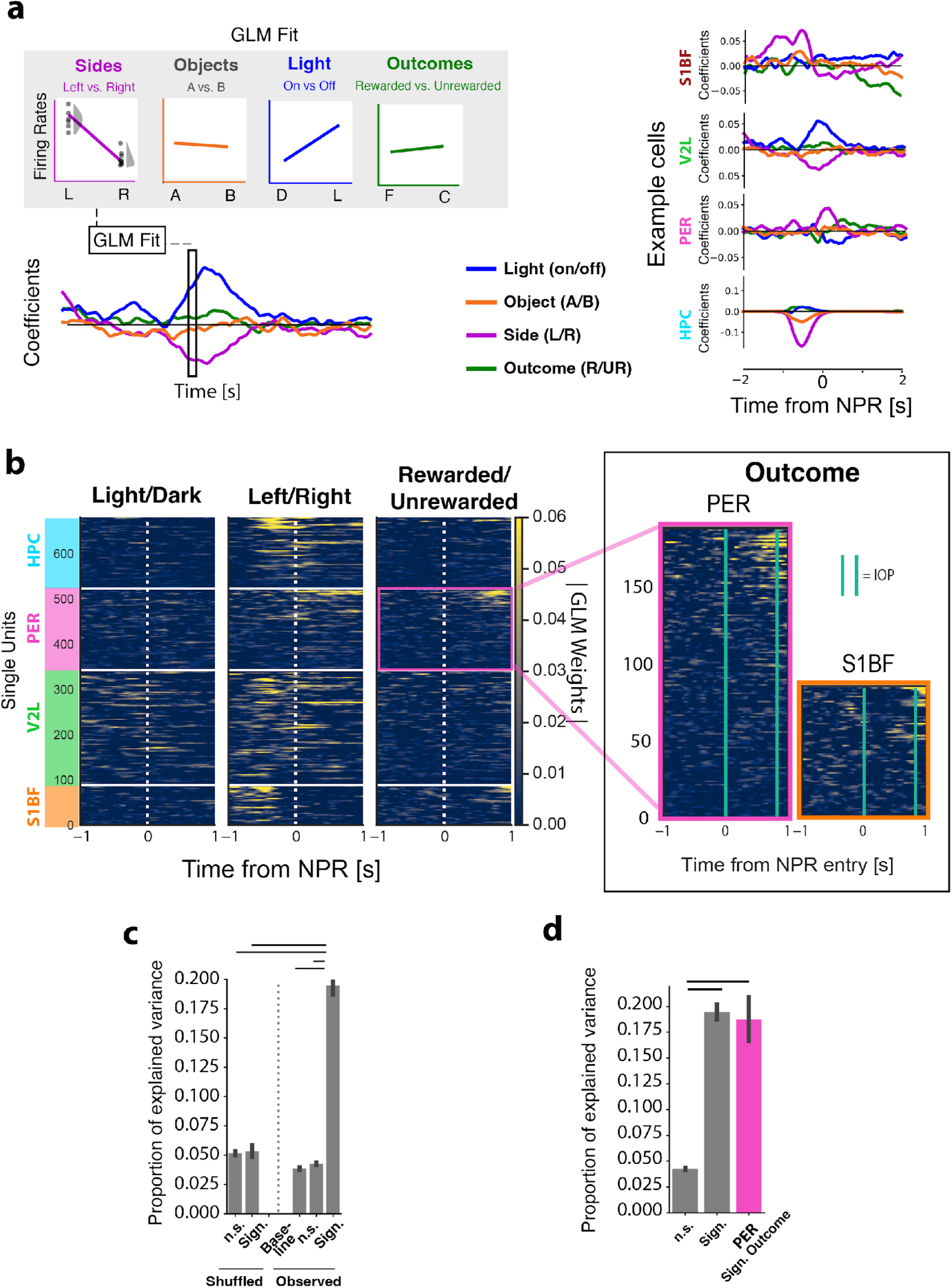
Generalized Linear Model of single unit encoding and population decoding of trial outcome. **a)** Left: Grey box is a schematic overview of Generalized Linear Model (GLM) fit at a given time point. Colored traces below the grey box are GLM coefficients quantifying neuronal encoding of photic condition, object identity, choice side and trial outcome over time by an example cell from V2L. R and UR are rewarded and unrewarded trials respectively. Right: GLM coefficients quantifying neuronal encoding of photic condition, object identity, choice side and trial outcome over time by example cells in S1BF, V2L, PER and HPC. NPR is Nose Poke for Reward. **b)** Left: GLM coefficient weight magnitude heatmap over time for different predictors (Light/Dark etc.), sorted per area and according to the timing of maximal outcome GLM weight magnitude. Right: close-up of Outcome coefficients over time for PER and S1BF, the two areas showing the clearest correlates in the Outcome period. **c)** GLM accuracy, scored by proportion of explained variance (R^2^ score) between predicted and actual firing rates. Model fits on time points that resulted in at least one significant predictor (Observed sign.) had significantly higher explained variance compared to fits on time points at which none of the predictors were significant (observed n.s.), as well as compared to time-points in a baseline period during the intertrial interval in which no relation was expected between firing rates and predictors. Model fits on time points that resulted in at least one significant predictor (Observed sign.) additionally had higher explained variance compared to chance-level model fits in which the trial labels were randomly shuffled (Shuffled: n.s. and Shuffled: Sign.). **d)** Verification of GLM outcome predictor accuracy in perirhinal cells. The explained variance from model fits on perirhinal cells that encoded the outcome (PER Sign. Outcome) was significantly higher than the explained variance from model fits that did not result in a significant predictor (n.s.).

## Notes

https://search.kg.ebrains.eu/instances/d406a98c-ae5c-4fb3-9f0c-4cf4de9b1094

### Competing Interest Statement

The authors have declared no competing interest.

